# The deubiquitylating enzyme Fat Facets promotes Fat signalling and restricts tissue growth

**DOI:** 10.1101/2024.01.27.577535

**Authors:** Lauren E. Dawson, Alexander D. Fulford, Aashika Sekar, Paulo S. Ribeiro

## Abstract

Tissue growth is regulated by many signals, including polarity cues. The Hippo signalling pathway restricts tissue growth and receives inputs from the planar cell polarity-controlling Fat signalling pathway. The atypical cadherin Fat restricts growth via several mechanisms that ultimately control the activity of the pro-growth transcriptional co-activator Yorkie. The Fat pathway modulates the activity of the Yorkie inhibitory kinase Warts, as well as the function of the FERM protein Expanded, which promotes Hippo signalling and also directly inhibits Yorkie. Although several Fat pathway activity modulators are known to be involved in ubiquitylation, the role of this post-translational modification in the pathway remains unclear. Moreover, no deubiquitylating enzymes have been described in this pathway. Here, using an *in vivo* RNAi screening approach, we identify the deubiquitylating enzyme Fat facets as a positive regulator of Fat signalling that is important for tissue growth control. Fat facets interacts genetically and physically with Fat signalling components and regulates transcription of Yorkie target genes. Thus, we uncover a role for reversible ubiquitylation in the control of Fat signalling and, by extension, in the regulation of tissue growth.

## Introduction

Developmental tissue growth and morphogenesis are controlled by a plethora of molecular mechanisms that must be tightly regulated to achieve reproducible organ and body size. In epithelia, one of the most important pathways involved in tissue growth regulation is the Hippo (Hpo) pathway, an evolutionarily conserved signalling cascade that integrates multiple signals that report on epithelial integrity ^**1-4**^. Hpo signalling culminates in the inhibition of Yorkie (Yki; mammalian YAP), a transcriptional co-activator that associates with transcription factors such as Scalloped (Sd; mammalian TEAD1-4) to promote the expression of genes involved in cell proliferation and inhibition of apoptosis ^**1, 5, 6**^. Yki activity is restrained by a kinase cascade consisting of the kinases Hpo and Warts (Wts; mammalian LATS1/2). The latter directly phosphorylates Yki, inhibiting its nuclear translocation, primarily by promoting interaction with 14-3-3 proteins ^**5, 7**^. Given its crucial role in tissue growth control, homeostasis is maintained by tight regulation of Hpo signalling, including a negative feedback loop in which Yki/YAP promote the expression of upstream activators of the kinase cascade ^**8–10**^.

Among the signals that regulate Hpo signalling are inputs from the cellular polarity machinery ^**3, 4, 9**^. Hpo signalling is regulated both by proteins involved in establishing and maintaining apico-basal polarity (e.g. Crumbs (Crb), Scribble (Scrib), among others) ^**4, 9**^, and planar cell polarity (PCP), such as the members of the Fat (Ft; mammalian FAT1-4) signalling pathway **^1^**^**1, 12**^. Ft is an atypical cadherin that localises to the sub-apical domain of epithelial cells and forms an heterotypic adhesion complex with the atypical cadherin Dachsous (Ds; mammalian DCHS1/2) ^**11**^, which is regulated by the Golgi resident kinase Four-jointed (Fj; mammalian FJX1) ^**13**^. The combination of the opposing Ds and Fj expression patterns in tissues, and the differential effect of Fj-mediated phosphorylation on the affinity of Ft and Ds to each other results in a gradient of Ft signalling that contributes to the regulation of PCP and tissue growth ^**11, 14–16**^. Ft-mediated regulation of growth involves several mechanisms. Ft inhibits the function of the atypical myosin Dachs (D) ^**17–19**^, which is a known negative regulator of Wts function, albeit the precise molecular mechanisms remain unclear ^**17, 20, 21**^. Ft also limits the activity of the zDHHC9-like transmembrane palmitoyltransferase Approximated (App) ^**22, 23**^ and the D-interacting protein Dachs ligand with SH3s (Dlish), which control D sub-cellular localisation and function ^**24, 25**^. Moreover, Dlish also inhibits Hpo signalling in a D-independent manner, via the regulation of the upstream activator Expanded (Ex) ^**26**^. Dlish interacts with Ex and promotes its degradation via the recruitment of Skp-Cullin-F-box (SCF) E3 ubiquitin ligase complexes containing the F-box protein Slimb, a known regulator of Ex function ^**27–29**^.

Interestingly, besides Dlish-mediated regulation of Ex stability, several steps of the Ft signalling pathway appear to be regulated by post-translational modifications such as ubiquitylation. D is thought to regulate Wts protein levels by an unknown mechanism that is likely to involve ubiquitylation ^**17, 20**^. In addition, Ft-mediated regulation of D function is at least partly dependent on the F-box protein Fbxl7, though it is still unclear whether D itself is ubiquitylated and degraded ^**30, 31**^. Finally, a recent report identified the E3 ligase Early girl (Elgi) as a new Ft signalling component involved in tissue growth regulation ^**32**^. Elgi is a D-interacting protein that controls D protein levels and, along with App, is proposed to control D and Dlish localisation to the apical membrane ^**32**^. Despite these observations, the precise molecular mechanisms by which ubiquitylation regulates Ft signalling and, by extension, tissue growth remain incompletely characterised. Importantly, to date there have been no reports of deubiquitylating enzymes (DUBs) as potential regulators of Ft signalling components. To address this, we performed an *in vivo* RNAi modifier screen to uncover DUBs involved in Ft signalling and identified Fat facets (Faf; mammalian USP9X) as a regulator of tissue growth. Faf genetically and physically interacts with Ft signalling components, regulates D sub-cellular localisation and controls expression of Yki target genes. Therefore, the function of Faf illustrates the crucial role of ubiquitylation in the regulation of Ft signalling and tissue growth.

## Results

### Identification of Faf as a deubiquitylating enzyme involved in Ft signalling

Previous studies have identified E3 ubiquitin ligases that are involved in the regulation of Ft signalling, such as Elgi and a Fbxl7-containing SCF complex ^**30–32**^. However, to date, a role for deubiquitylating enzymes (DUBs) in the regulation of Ft signalling has not been described. To identify DUBs that modulate Ft function, we performed an *in vivo* RNAi modifier screen using the *Drosophila* adult wing as a model **(Figure S1A)**. We used the wing driver *nub-Gal4* (*nub>*) to target the full complement of *Drosophila* DUBs with *UAS-RNAi* transgenes. Each *DUB^RNAi^* line was crossed to *nub-Gal4, UAS-ft* (*nub>ft*) or *nub-Gal4* (*nub>*) as a control. **Figures S1B** and **S1C** show the results of the *in vivo* screens and the quantification of the relative wing size of the different genotypes tested. In agreement with its effect on Hpo and PCP signalling, *UAS-ft* expression in the wing pouch resulted in smaller and rounder wings **(Figure 1D, S1C and S2C-E)** compared with controls **(Figure 1A, S1D, S2A and S2C-E)**. As a result of our screening approach, we identified the DUB Fat facets (Faf, encoded by *faf*, *CG1945*) as a potential regulator of Ft signalling. *faf* depletion using several independent RNAi lines resulted in a partial suppression of the Ft undergrowth phenotype **(Figure 1E, 1J and S1C)**. Interestingly, Faf seems to primarily affect the tissue growth function of Ft, but not its PCP function, based on the ratio of the lengths of the anterior-posterior (AP) and proximal-distal (PD) axes of the adult wing and wing circularity (**Figure S2B-E**; see Materials and Methods for details). Next, we sought to determine if modulation of *faf* expression alone affects tissue growth and to validate its interaction with Ft signalling. *faf* depletion in the developing wing using *nub-Gal4* resulted in a mild increase in wing size, when compared to controls **(Figure 1J and S1E-G)**. Conversely, over-expression of *faf* (Faf^isoform^ ^C^ ^**33**^; **Figure 2A**) reduced wing size **(Figure 1C, 1J and S1H)**. When combined with *UAS-ft*, depletion of *faf* suppressed the Ft phenotype **(Figure 1E and 1J)**, while *faf* over-expression enhanced it **(Figure 1F and 1J)**. In contrast, simultaneous depletion of *ft* and *faf* caused an enhancement of the *ft^RNAi^* phenotype **(Figure 1G, 1H and 1J)**, while over-expression of *faf* in the context of *ft^RNAi^* partially suppressed the wing overgrowth phenotype **(Figure 1G, 1I and 1J)**. Interestingly, we observed that modulation of Faf levels resulted in a very specific defect of the L2 wing vein, with the appearance of extra wing material in *faf^RNAi^*wings **(Figure 1B, S2H and S2I)**. Notably, this phenotype has previously been associated with changes in Ft levels and was also observed when Ft levels were modulated in isolation **(Figure 1D and 1G)** ^**34, 35**^. Importantly, modulating Faf levels enhanced the L2 wing vein phenotypes **(Figure S2I)**. Our results suggest that Faf promotes Ft activity in tissue growth control.

**Figure 1.**
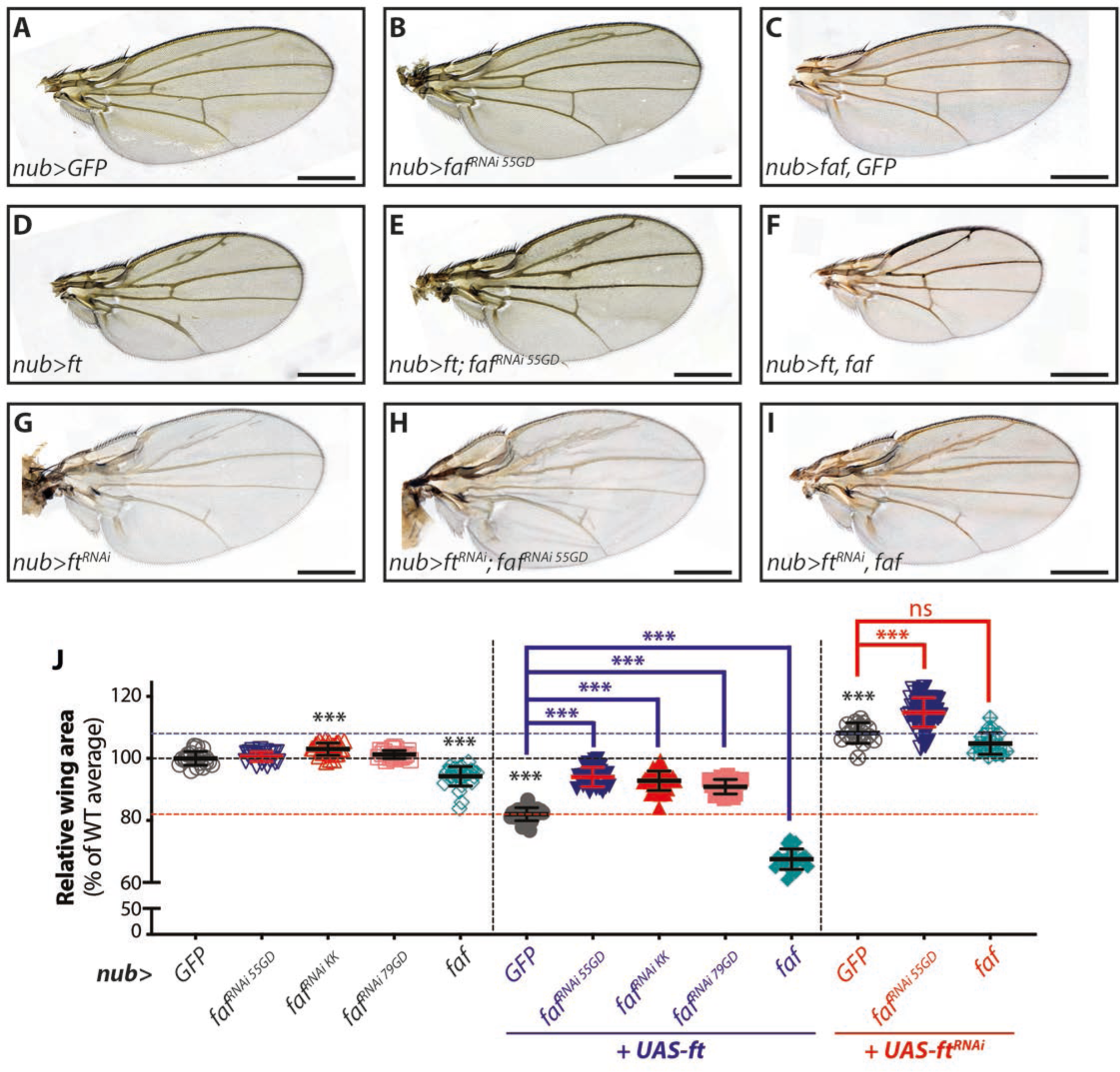
Faf modulates Ft-mediated regulation of tissue growth. **(A-I)** Modulation of Faf expression affects tissue growth during normal development or in conditions when Ft is over-expressed or depleted. Shown are adult wings from flies raised at 25°C expressing the indicated transgenes in the wing pouch under the control of *nub-Gal4* (*nub>*). Compared to control adult wings expressing GFP (A), Ft-expressing wings were smaller (D), while *ft* depletion caused increased growth (G). Depletion of *faf* mildly enhanced tissue growth (B), while *faf* over-expression resulted in undergrowth (C). *faf* depletion resulted in a partial rescue of the undergrowth phenotype caused by *UAS-ft* (E), while it enhanced the overgrowth phenotype of *ft^RNAi^* flies (H). In contrast, *UAS-faf* enhanced the growth impairment of *UAS-ft* flies (F) and mildly suppressed the overgrowth phenotype of *ft^RNAi^* flies (I). **(J)** Quantification of relative adult wing sizes from flies expressing the indicated transgenes under the control of *nub-Gal4*. Data are represented as % of the average wing area of the respective controls (*nub>GFP*, average set to 100%). Data are shown as average ± standard deviation, with all data points depicted. n>19 for all genotypes. Significance was assessed using Brown-Forsythe and Welch ANOVA analysis comparing all genotypes to their respective controls (*nub>GFP*, *nub>ft* or *nub>ft^RNAi^*; black, blue or red asterisks, respectively) with Dunnett’s multiple comparisons test. *, p<0.05; **, p<0.01; ***, p<0.001; ns, non-significant. Scale bar represents 500 μm.

**Figure 2.**
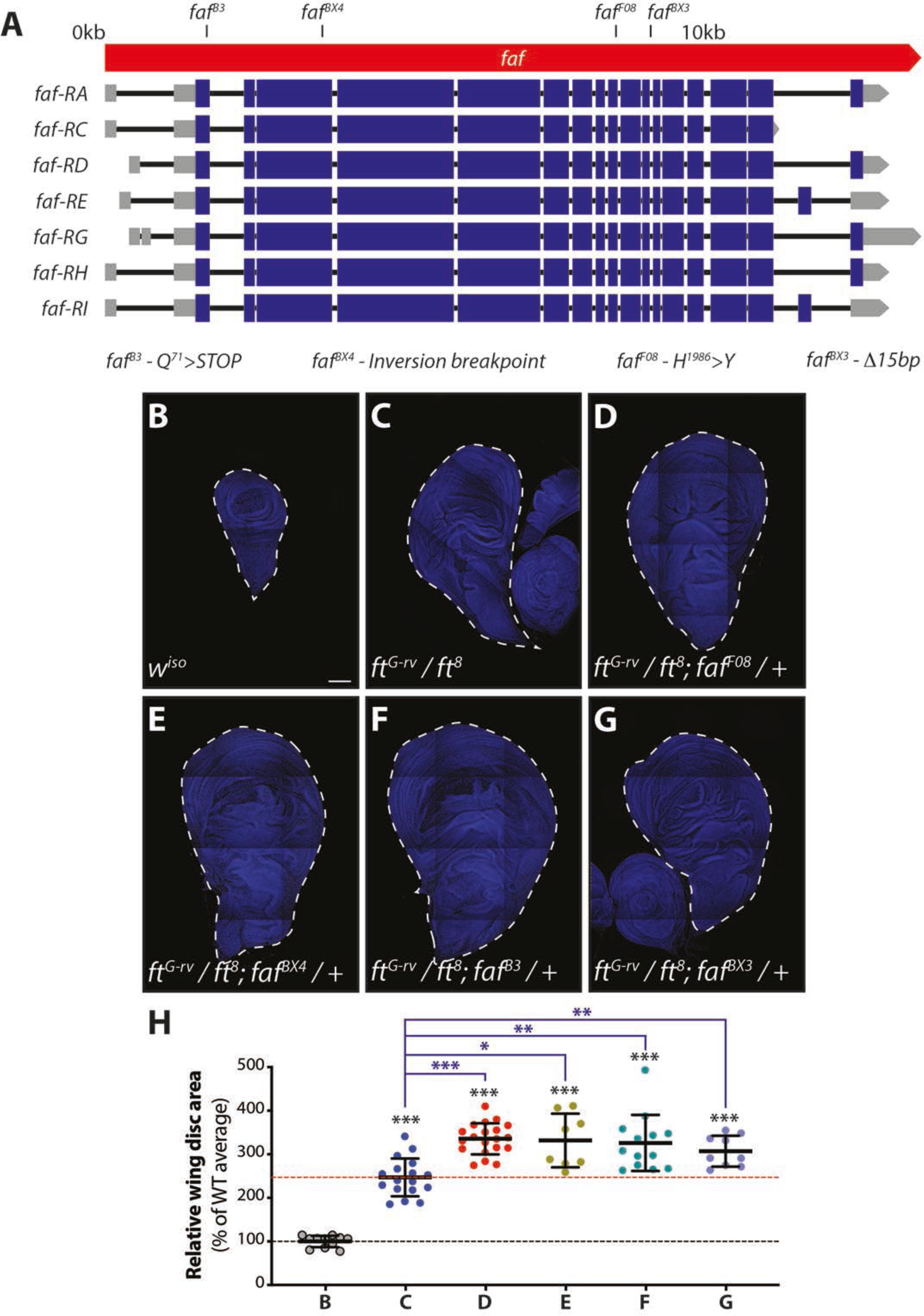
*faf* genetically interacts with *ft*. **(A)** Schematic representation of the *faf* locus. mRNA transcripts encoded in the *faf* locus are represented (boxes depict exons, while introns are shown as black lines. Coding sequences are shown as blue boxes, and UTRs are represented as grey boxes). Mutant alleles used are mapped onto the *faf* locus. *faf ^B3^* corresponds to a nonsense mutation leading to a truncated Faf protein (Q^71^>STOP). The *faf ^BX4^* lesion is the site of an inversion breakpoint. *faf ^F08^* is a point mutation in the DUB catalytic domain (H^1986^>Y). *faf ^BX3^* corresponds to a 15bp deletion in the vicinity of an exon-intron boundary. **(B-G)** Loss of one copy of *faf* enhances the overgrowth of *ft* mutants. Shown are tiled confocal images of wing imaginal discs. Compared to control L3 imaginal discs (*w^iso^*, B), the transheterozygous combination of *ft^G-rv^* and *ft^8^* leads to a dramatic increase in tissue size (C), which is further enhanced by the presence of a *faf* mutant allele (*faf ^F08^* (D), *faf ^BX4^* (E), *faf ^B3^* (F), *faf ^BX3^* (G)). **(H)** Quantification of relative wing imaginal disc size. Data are represented as % of the average imaginal wing disc area of controls (*w^iso^*, which was set to 100% and indicated by the black dashed line). Red dashed line indicates average wing disc size of *ft* transheterozygous mutants. Data are shown as average ± standard deviation, with all data points depicted. n=11, 17, 20, 8, 13 and 9 for B, C, D, E, F and G, respectively. Significance was assessed using Brown-Forsythe and Welch ANOVA analyses comparing all genotypes to the respective control (*w^iso^* or *ft^G-rv^/ft^8^*; black or blue asterisks, respectively), with Dunnett’s multiple comparisons test. *, p<0.05; **, p<0.01; ***, p<0.001. Scale bar represents 100 μm.

To validate the results obtained with *faf^RNAi^*, we next assessed if *faf* genetically interacts with *ft* mutations in the regulation of tissue growth. Due to its critical role in controlling Hpo signalling, *ft* homozygous mutations are lethal and associated with extreme tissue overgrowth phenotypes ^**36, 37**^. However, certain *ft* mutations allow the analysis of tissues from late L3 larvae in a trans-heterozygous situation, such as the *ft^G-rv^*/*ft^8^*combination, allowing them to be studied alongside other genetic alterations ^**15, 38**^. *ft^G-rv^*/*ft^8^*trans-heterozygous mutant wing discs displayed extreme tissue overgrowth, compared to wild-type (*w^iso^*) wing discs (∼170% larger than controls; compare **Figure 2B and 2C**). Remarkably, when we combined *ft^G-rv^*/*ft^8^* mutations with various *faf* mutant alleles **(Figure 2A)**, we observed an enhancement of the tissue overgrowth phenotype **(Figure 2D-H)**. This indicates that *faf* genetically interacts with *ft* and that Faf function is important to restrict tissue growth. Given that *faf* mutation enhances *ft* mutant phenotypes, it is possible that Faf controls growth in both Ft-dependent and Ft-independent manners. Alternatively, Faf may act on residual Ft protein in *ft^G-rv^*/*ft^8^* trans-heterozygotes. Taken together, our data in the *Drosophila* wing are consistent with Faf promoting the tissue growth suppressing function of Ft.

### Faf genetically interacts with core Ft signalling proteins

Having observed that Faf genetically interacts with Ft and plays an important role in the regulation of tissue growth, we extended our analysis to other members of the Ft signalling pathway that directly interact with Ft, such as Ds, Dlish and Fbxl7 **(Figure 3)**. Ft and Ds regulate tissue growth at least in part via their physical interaction across cell boundaries ^**11**^. This interaction enhances the activity of Ft, thereby promoting its growth-suppressing function ^**11**^. Accordingly, we found that Ds over-expression caused a reduction in wing size **(Figure 3B and 3P)**. Similarly to what was observed with Ft, *faf^RNAi^* reversed this Ds-induced phenotype **(Figure 3C-E and 3P)**. This is consistent with a positive role for Faf in Ft signalling. We also assessed if Faf could modulate phenotypes caused by downstream effectors of Ft signalling, such as Dlish and Fbxl7 ^**24, 25, 30, 31**^. Dlish negatively regulates Hippo signalling and, therefore, promotes tissue growth ^**24–26**^. Accordingly, depletion of Dlish (*Dlish^RNAi^*) in the developing wing resulted in reduced tissue growth **(Figure 3G and 3Q)**. Co-depletion of *faf* and *Dlish* suppressed this phenotype **(Figure 3H and 3Q)**, whilst Faf over-expression enhanced the undergrowth **(Figure 3I and 3Q)**. As Dlish function is influenced by Ft, we combined *Dlish^RNAi^* with Ft over-expression, which resulted in an enhancement of the *Dlish^RNAi^*-induced undergrowth **(Figure 3J and 3Q)**. To address this is dependent on Faf activity, we co-depleted *faf* in these conditions (*Dlish^RNAi^* + *ft* over-expression), which led to a rescue of the phenotype **(Figure 3K and 3Q)**, suggesting that Faf does indeed modulate Dlish phenotypes via Ft. As previously observed, *faf^RNAi^* mostly affected wing size rather than shape, as wings remained rounder than controls **(Figure 3K)**, further indicating that Faf has a minor role in Ft-mediated regulation of tissue shape. In agreement with its previously reported role, over-expression of Fbxl7 leads to a significant reduction in tissue size ^**30, 31**^ **(Figure 3L and 3R)**. Interestingly, depletion of *faf* did not significantly affect the Fbxl7 phenotype, suggesting that Faf may act at the level of Fbxl7 **(Figure 3M and 3R)**. Moreover, Ft over-expression failed to enhance the Fbxl7 undergrowth phenotype **(Figure 3N and 3R)**, which is consistent with Fbxl7 playing a crucial role downstream of Ft. *faf^RNAi^* only had a slight effect on tissue growth in these conditions **(Figure 3O and 3R)**. We then extended our observations to additional Ft-associated proteins, such as Ex, Elgi and App **(Figure S3)**. Our results suggest that Faf modifies phenotypes associated with Ex **(Figure S3B-E and S3P)** and Elgi **(Figure S3F-J and S3Q)**, but not with App **(Figure S3K-O and S3R)**. Taken together, these results suggest that Faf genetically interacts with multiple Ft signalling components and may work at the level of Fbxl7 and App, and potentially antagonistically to Dlish.

**Figure 3.**
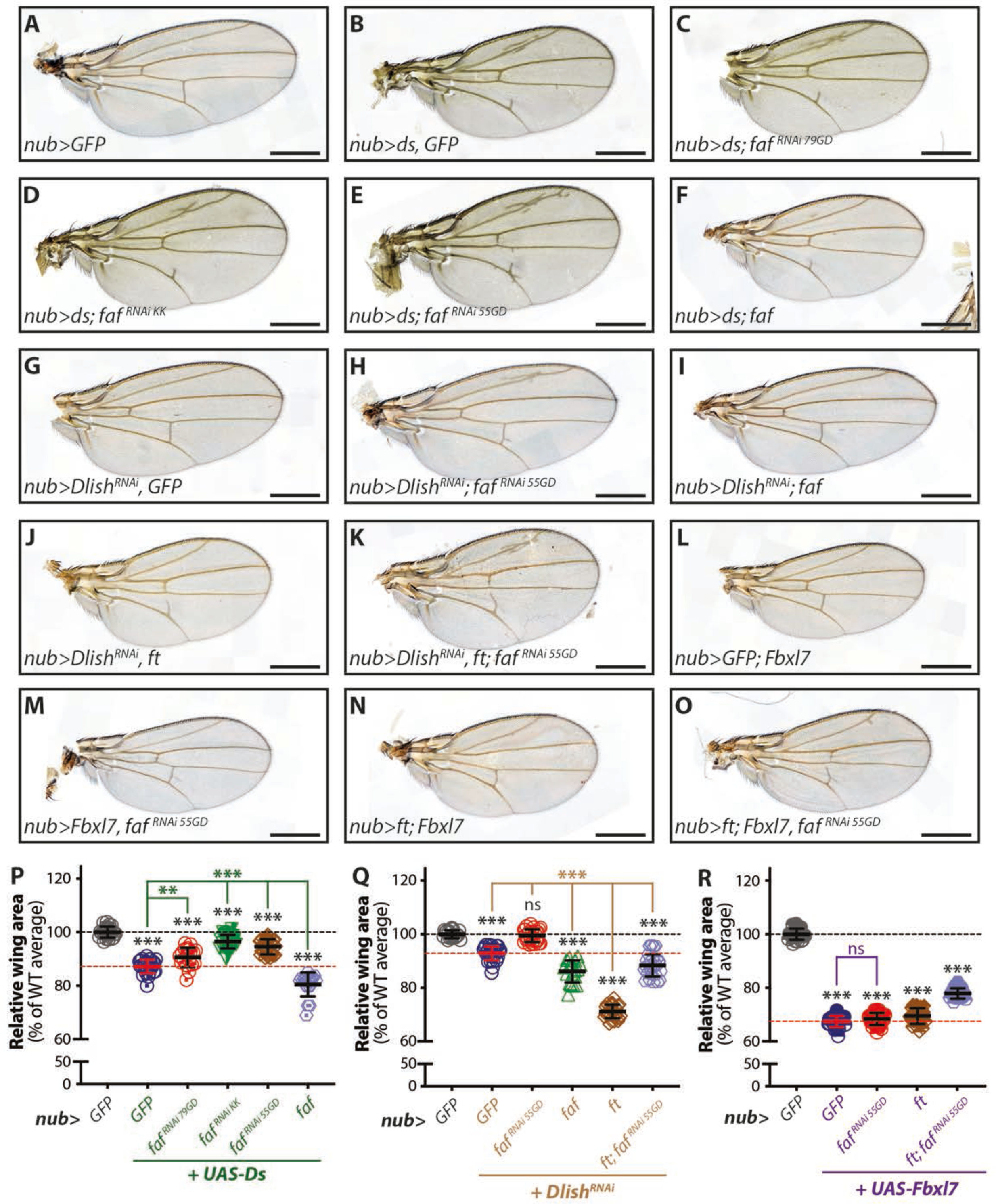
*faf* genetically interacts with *ds*, *Dlish* and *Fbxl7* in the regulation of wing tissue growth. **(A-O)** Modulation of Faf expression levels affects the tissue growth phenotypes of *ds-* (B-F), *Dlish^RNAi^*-(G-K) and *Fbxl7*-expressing flies (L-O). Shown are adult wings from flies raised at 25°C expressing the indicated transgenes in the wing pouch under the control of *nub-Gal4* (*nub>*). Compared to control adult wings (A), wings expressing *ds* displayed significant tissue undergrowth (B), which was partially rescued by simultaneous depletion of *faf* (C, D and E) and enhanced by *faf* over-expression (F). RNAi-mediated depletion of *Dlish* (G) caused a mild reduction in wing size, which was partially suppressed by co-depletion of *faf* (H), and enhanced by *faf* over-expression (I). Expression of *ft* resulted in an enhancement of the undergrowth of *Dlish^RNAi^* wings (J), and this was in part prevented by *faf^RNAi^*(K). *Fbxl7* over-expression caused reduced wing size compared to controls (L) and this was not significantly affected by *faf^RNAi^* (M) or *ft* expression (N), but this was partially suppressed when *ft* was combined with *faf^RNAi^* (O). **(P-R)** Quantification of relative adult wing sizes in genetic interactions with *ds* (P), *Dlish* (Q) or *Fbxl7* (R). Data are represented as % of the average wing area of control wings (*nub>GFP*, which were set as 100%). Data are shown as average ± standard deviation, with all data points represented. n>18 for all genotypes. Black dashed lines represent average size of controls (100%) whilst red dashed lines indicate average size of adult wings from flies expressing *ds*, *Dlish^RNAi^* or *Fbxl7* under the control of *nub-Gal4*. Significance was assessed using a one-way ANOVA comparing all genotypes to their respective controls (*nub>GFP*, *nub>GFP*+*UAS-Ds*, *nub>GFP*+*Dlish^RNAi^*or *nub>GFP*+*UAS-Fbxl7*; black, green, brown or blue asterisks, respectively), with Dunnett’s multiple comparisons test. **, p<0.01; ***, p<0.001. n.s. non-significant. Scale bar represents 500 μm.

### Faf physically interacts with Ft and regulates its protein levels

Next, we assessed whether the genetic interactions between *faf* and members of the Ft signalling pathway could be the result of specific protein-protein interactions. For this, we expressed Faf in *Drosophila* S2 cells (**Figure 4A**; Faf^LD^, a protein encoded by the cDNA clone *LD22582*) and performed co-immunoprecipitation (co-IP) assays with the intracellular region of Ft (Ft^ICD^ ^**39**^). Ft^ICD^ was readily detected in Faf^LD^ co-IPs, but not in the respective GFP controls **(Figure 4B and S4A)**. In agreement with the widespread role of DUBs as regulators of protein stability, we noticed that the protein levels of Ft^ICD^ in cell lysates appeared higher when Ft^ICD^ was co-expressed with Faf^LD^ **(Figure 4B)**. Therefore, we conducted further experiments using Faf^LD^ expression or *faf* RNAi-mediated depletion to validate this observation. Indeed, Ft^ICD^ was stabilised in the presence of Faf^LD^ **(Figure 4C and S4B)** and, conversely, Ft^ICD^ levels were reduced when endogenous *faf* was depleted from S2 cells **(Figure 4D, S4C and S4D)**. This suggests that Faf may regulate Hippo signalling and tissue growth by modulating Ft protein levels.

**Figure 4.**
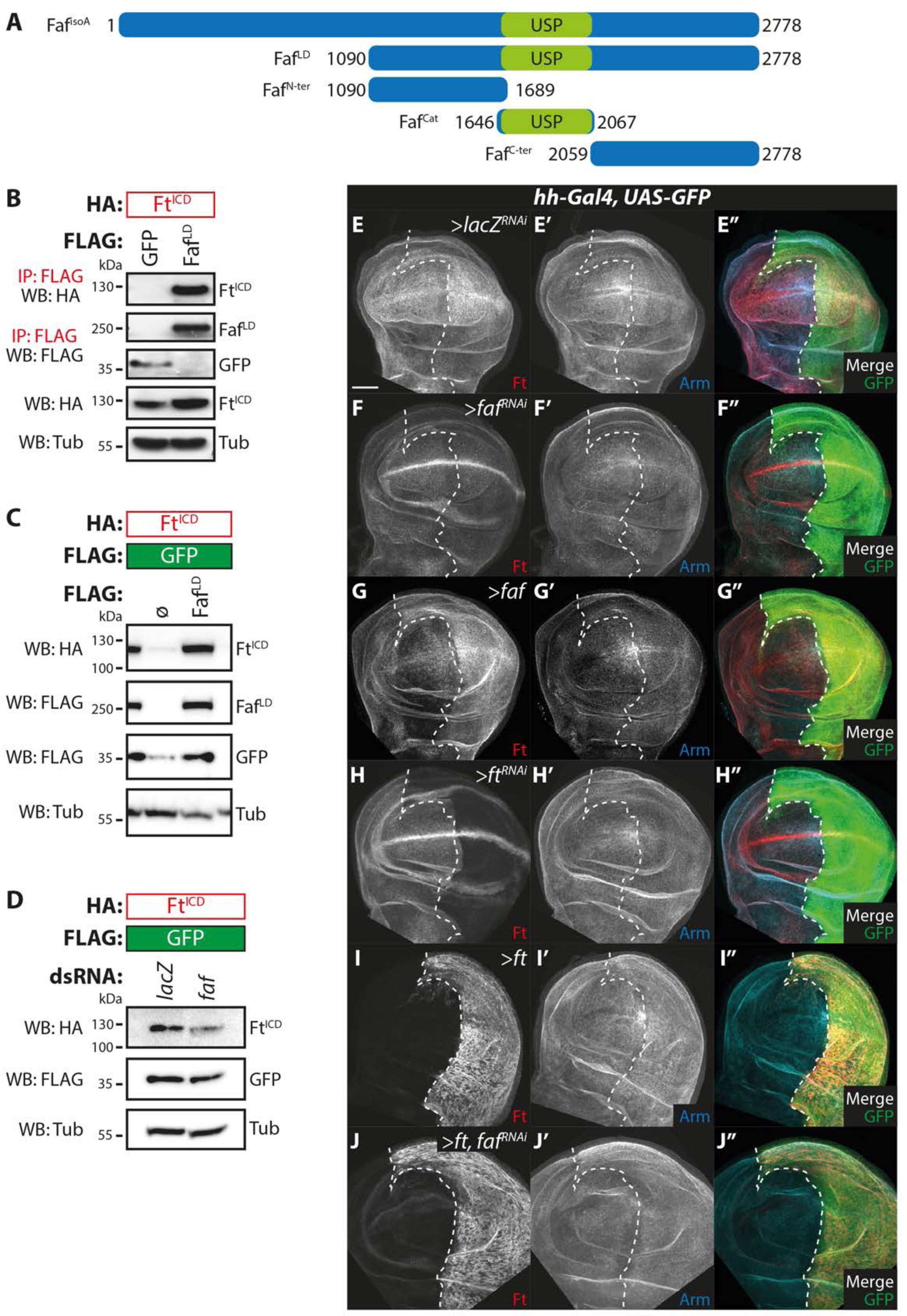
Faf interacts with Ft and regulates its protein levels *in vitro* and *in vivo*. **(A)** Schematic representation of the Faf constructs used in this study. Numbers denote amino acid position. USP DUB domain is represented in green. **(B)** Faf binds to Ft. HA-tagged Ft^ICD^ was co-expressed with FLAG-tagged GFP or Faf^LD^ (B) in *Drosophila* S2 cells. Cells were lysed and lysates were subjected to co-immunoprecipitation using FLAG agarose beads. Lysates were analysed by immunoblot using the indicated antibodies for detection of protein expression and co-purification. Tubulin (Tub) was used as loading control. **(C, D)** Faf regulates Ft protein levels in *Drosophila* S2 cells. (C) HA-tagged Ft^ICD^ was expressed in S2 cells in the presence or absence of Faf^LD^. 48h after transfection, cells were lysed and lysates were analysed by immunoblot using the indicated antibodies. Ø represents expression of empty vector. (D) S2 cells were treated with the indicated dsRNAs 24h before co-transfection with the indicated constructs. Ft^ICD^ protein levels were analysed by Western blotting with the indicated antibodies 48h after cell transfection. GFP and Tubulin (Tub) were used as transfection and loading control, respectively. **(E-J)** Faf regulates Ft protein levels *in vivo*. Shown are XY confocal micrographs of third instar wing imaginal discs expressing the indicated constructs under the control of *hh-Gal4*, showing Ft antibody staining (E-J and red in E’’-J’’), Arm antibody staining (E’-J’ and cyan in E’’-J’’), and direct fluorescence from GFP (green in E’’-J’’ merged images). Compared to the controls (E, *lacZ^RNAi^*), *faf* depletion (F) and *faf* over-expression (G) resulted in a decrease or increase in Ft protein levels, respectively. Shown are also *ft^RNAi^* (H) and *ft* over-expression (I) controls to validate antibody specificity. Depletion of *faf* in the presence of *ft* over-expression resulted in a mild decrease in Ft protein levels (J). Ventral is up in XY sections, whilst GFP marks the *hh-Gal4*-expressing posterior compartment (right). Dashed white line depicts boundary between anterior and posterior compartments. Scale bar represents 50 μm.

To validate these observations *in vivo*, we modulated Ft or Faf expression in the posterior compartment of the wing disc (using *hh-Gal4*) and assessed the effect on Ft protein levels using a specific antibody ^**19, 39, 40**^ **(Figure 4H)**. As a control for general effects on the levels of proteins localised at the apical cell surface, we monitored Armadillo (Arm; *Drosophila* β-catenin) protein levels. Altering Faf levels *in vivo* recapitulated the results observed in S2 cells, and *faf^RNAi^*expression resulted in a reduction in Ft levels **(Figure 4E and 4F)**, whilst over-expression of Faf increased Ft protein levels **(Figure 4E and 4G)**. Importantly, the effect of Faf on Ft levels was specific as Arm levels were generally unaffected **(Figure 4, Figure S4E-G)**. Together, these data suggest that Faf regulates Ft protein levels both in *Drosophila* S2 cells and *in vivo*.

### Faf modulates Dachs subcellular localisation

Next, we tested whether Faf regulates events downstream of Ft and focused on a potential modulation of D function. As extensively documented, D is one of the most important effectors of Ft signalling ^**15, 17, 18**^, and Ft regulates D by controlling its subcellular localisation ^**18**^. To assess D localisation, we used a *D::GFP* knock-in allele in combination with *en-Gal4, UAS-RFP*. This allowed us to monitor D subcellular localisation in the posterior compartment of the developing wing disc (marked by RFP) and to use the anterior compartment as an internal control **(Figure 5A)**. For each experimental condition, we imaged the control and the posterior compartments, using Arm as a control for global effects on apical protein localisation. As seen in **Figure 5B**, in control flies (*lacZ^RNAi^*), D is localised at the membrane, similarly to Arm, and the stereotypical D polarisation toward the distal side of the cell can be observed. In agreement with published data ^**18, 20, 22, 30, 31, 41, 42**^, over-expression of Ft resulted in the mislocalisation of D, which appeared both more cytoplasmic and less polarised at the membrane, compared to controls **(Figure 5C)**. In agreement with our previous results, we observed that combining Ft over-expression with depletion of *faf* (*UAS-ft, UAS-faf^RNAi^*) resulted in a return to control conditions **(Figure 5D)**. In contrast, when we tested the effect of *faf* over-expression, we observed loss of D polarisation, increased levels of D in the cytoplasm and, in some cases, patches of tissue where D was not localised at the membrane, a phenotype similar to Ft over-expression ^**18**^. Depletion of *ft* or *faf* resulted in a less apparent D polarisation when compared to the anterior control compartment **(Figure S5C and S5D)**. Together, these results suggest that Faf is, at least partly, required for Ft-mediated regulation of D polarisation and that modulating Faf levels is sufficient to alter the pattern of D in tissues.

**Figure 5.**
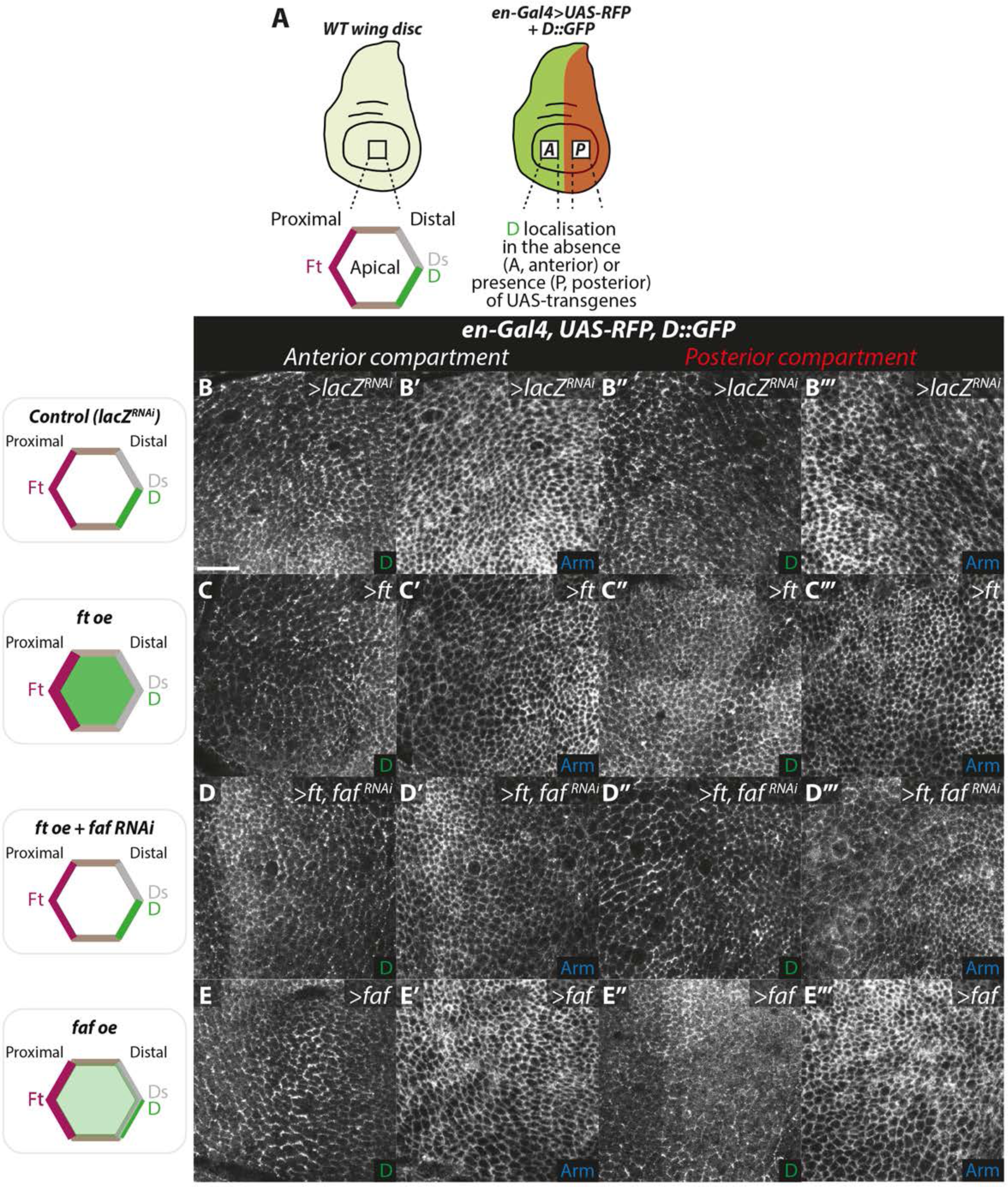
Ft regulates subcellular localisation of D in a Faf-dependent manner. **(A)** Schematic representation of subcellular localisation of Ft signalling components and experimental setting. D localisation was monitored using a D::GFP transgene, whilst expression of genes of interest was controlled by *en-Gal4*, which is expressed in the posterior compartment of the wing imaginal disc (marked by RFP expression). **(B-E)** Regulation of D subcellular localisation by Ft and Faf. XY sections of third instar wing imaginal discs, showing direct fluorescence from GFP in the control anterior compartment (B-E) or experimental posterior compartment (B’’-E’’), or Arm staining in the anterior (B’-E’) or posterior (B’’’-E’’’) compartments. Cartoons represent expected results for each genotype tested. Over-expression of *ft* or *faf* results in mislocalisation of D, which appears more cytoplasmic (C’’ and E’’), compared to controls (B’’). The *ft* over-expression phenotype was largely rescued by co-depletion of *faf* (D’’). No overt effects were observed for Arm in all genotypes tested (B’’’-E’’’). Scale bar represents 10 μm.

### Faf regulates Yki target genes in vivo

Our data suggest that Faf regulates tissue growth by modulating the function of Ft. To confirm that the effects of Faf on tissue growth were indeed due to changes in signalling activity downstream of Ft, we tested whether Faf could affect the expression of genes known to be regulated by Yki, the main effector protein regulated by the Hippo pathway. We monitored Yki-mediated transcription using two widely used Hippo signalling *in vivo* reporters; *ex-lacZ* ^**43**^ **(Figure 6A-F)** and *HRE-diap1::GFP* ^**44**^ **(Figure 6G-L)**. Transgenes were specifically expressed in the posterior compartment of the wing using *en-Gal4* as a driver and marked by the expression of GFP or RFP, respectively in the *ex-lacZ* or *DIAP1::GFP* experiments. As a control, we used *hpo^RNAi^*, which is known to lead to increased Yki-mediated gene expression ^**7, 45**^ **(Figure 6B, 6F, 6H and 6L)**. In agreement with our observations regarding the effect of Faf on tissue growth, depletion of *faf* (*faf^RNAi^*) caused an increase in Yki-mediated transcription, consistent with decreased Hippo signalling activity **(Figure 6C, 6F, 6I and 6L)**. Accordingly, *faf* over-expression resulted in decreased Yki activity **(Figure 7A, 7B and 7E)**. We also used the Yki-mediated transcriptional readout to determine whether Faf is acting through Ft to produce these effects. Over-expression of Ft in the posterior compartment of the wing resulted in a significant decrease in Yki-mediated transcription, consistent with its role in activating Hippo signalling **(Figure 6D, 6F, 6J and 6L)**. Interestingly, depletion of *faf* completely abrogated the effect of Ft over-expression on the expression of Yki target genes **(Figure 6E, 6F, 6K and 6L)**. Indeed, the expression levels of the *ex-lacZ* and *DIAP1::GFP* reporters were significantly different in the *UAS-ft, UAS-faf^RNAi^* condition compared to when Ft was overexpressed in isolation, and were closer to levels seen in controls. Together, this data suggests that Faf promotes tissue growth by positively regulating Hippo signalling and that it does so by modulating the function of Ft.

**Figure 6.**
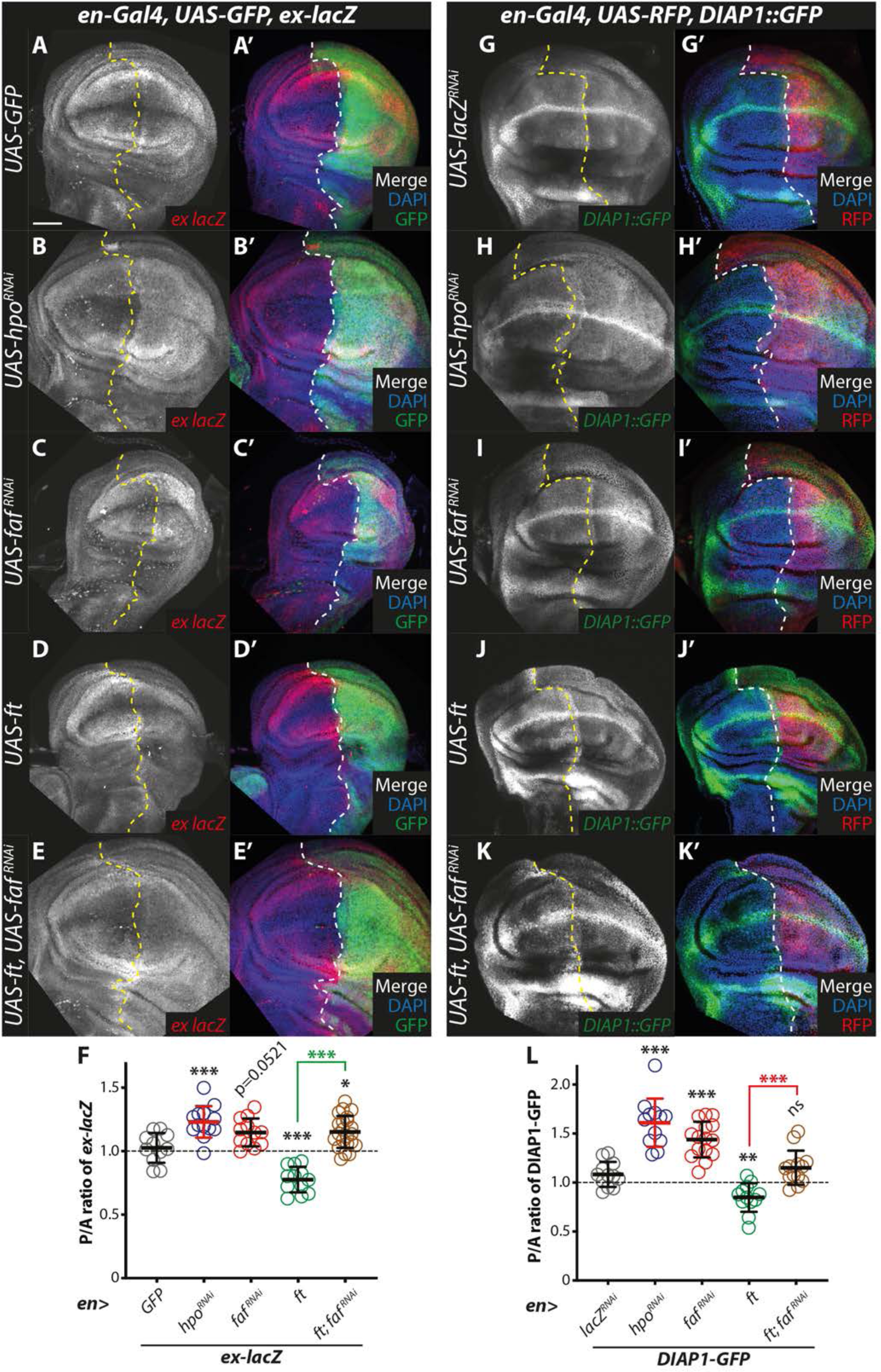
Faf regulates expression of Yki target genes *in vivo*. **(A-E)** Regulation of *ex-lacZ* expression. XY confocal sections of third instar wing imaginal discs containing *ex-lacZ* (A-E, shown in red in A’-E’ merged images), in which *en-Gal4* was used to drive expression of *UAS-GFP* (A), *UAS-hpo^RNAi^*(B), *UAS-faf^RNAi^*(C), *UAS-ft* (D) or *UAS-ft* and *UAS-faf^RNAi^* (E). Compared to controls (A), *hpo* (B) or *faf* (C) depletion resulted in an increase in *ex-lacZ* levels in the posterior compartment, indicating lower Hpo pathway activity and, consequently, higher levels of Yki-mediated transcription. As expected, over-expression of *ft* resulted in decreased *ex-lacZ* expression (D), which was abrogated by simultaneous depletion of *faf* (E). GFP (green in A’-E’ merged images) indicates posterior compartment where transgenes are expressed. DAPI (blue) stains nuclei. Dashed lines indicate boundary between anterior and posterior compartments. **(F)** Quantification of *ex-lacZ* expression levels. Shown are the posterior/anterior (P/A) *ex-lacZ* ratios for the different genotypes analysed. Data are shown as average ± standard deviation, with all data points represented. n>9 for all genotypes. Significance was assessed using a one-way ANOVA comparing all genotypes to their respective controls, with Dunnett’s multiple comparisons test. **(G-K)** Regulation of *DIAP1::GFP* expression. XY confocal sections of third instar wing imaginal discs carrying *DIAP1::GFP* (G-K, shown in green in G’-K’ merged images), in which *en-Gal4* was used to drive expression of *UAS-lacZ^RNAi^* (G), *UAS-hpo^RNAi^* (H), *UAS-faf^RNAi^*(I), *UAS-ft* (J) or *UAS-ft* and *UAS-faf^RNAi^*(K). Compared to controls (G), *hpo* (H) or *faf* (I) depletion resulted in an increase in *DIAP1::GFP* levels in the posterior compartment. As expected, *ft* over-expression resulted in decreased *DIAP1::GFP* levels (J), which was suppressed by simultaneous depletion of *faf* (K). RFP (red in G’-K’ merged images) indicates posterior compartment where transgenes are expressed. DAPI (blue) stains nuclei. Dashed lines indicate boundary between anterior and posterior compartments. **(L)** Quantification of *DIAP1::GFP* expression levels. Shown are the posterior/anterior (P/A) *DIAP1::GFP* ratios for the different genotypes analysed. Data are shown as average ± standard deviation, with all data points represented. n>8 for all genotypes. Significance was assessed using a one-way ANOVA comparing all genotypes to their respective controls (*en>GFP* or *UAS-ft*; black or green and red asterisks, respectively), with Dunnett’s multiple comparisons test. *, p<0.05; **, p<0.01; ***, p<0.001; ns, non-significant. Scale bar represents 50 μm.

**Figure 7.**
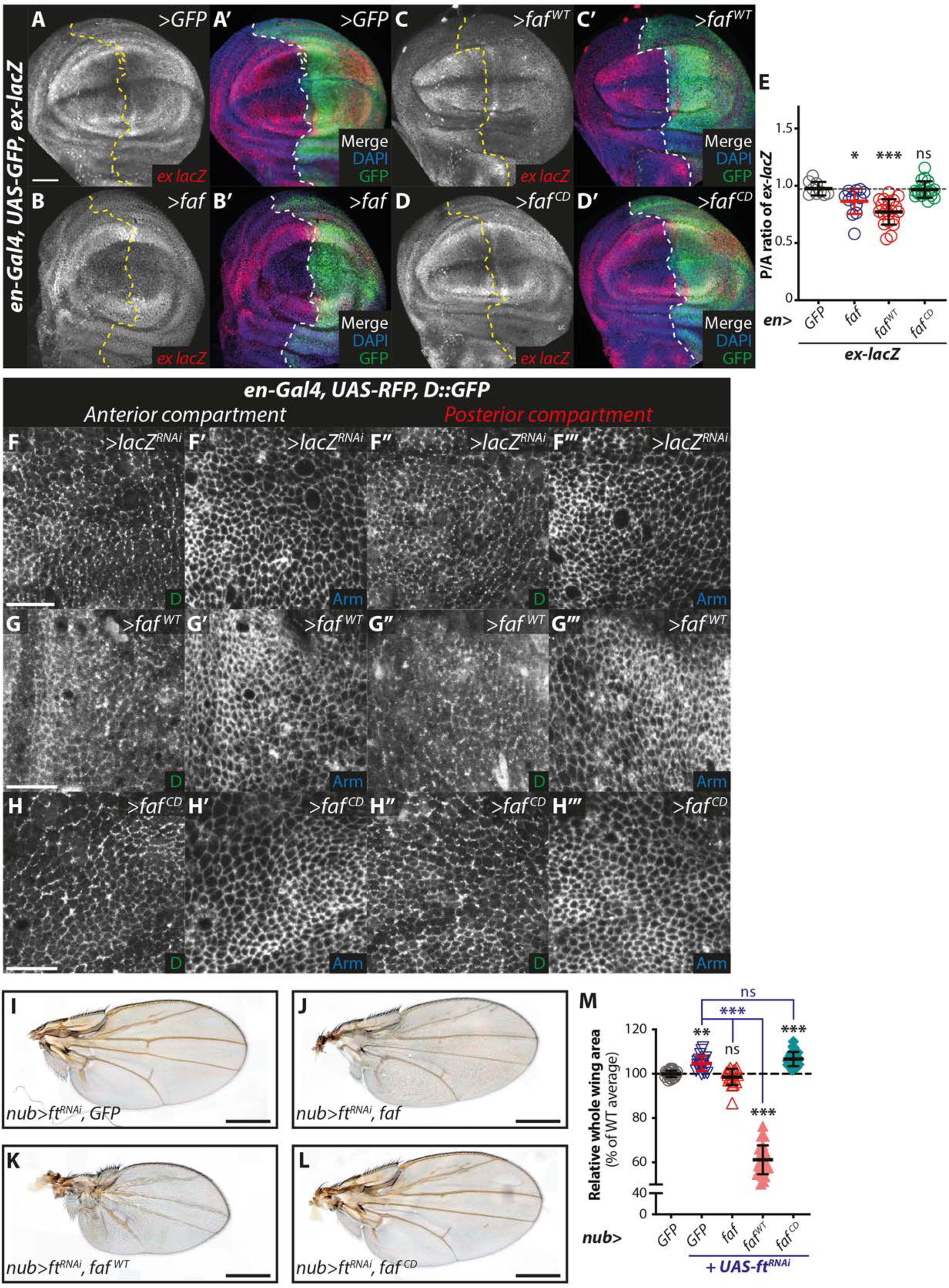
The role of Faf in modulating Ft function is dependent on its DUB activity. **(A-D)** Faf regulates Yki target gene expression in a DUB-dependent manner. XY confocal sections of third instar wing imaginal discs carrying *ex-lacZ* (A-D, shown in red in A’-D’ merged images), in which *en-Gal4* was used to drive expression of *GFP* (A), *faf* (B), *faf^WT^* (C) or *faf^CD^*(D). Compared to controls (A), *faf* over-expression (B,C) resulted in a decrease in the P/A ratio of *ex-lacZ*, indicating lower levels of Yki-mediated transcription. Expression of a Faf catalytic mutant had no effect (D). GFP (green in A’-D’ merged images) indicates posterior compartment where transgenes are expressed. DAPI (blue) stains nuclei. Dashed lines indicate boundary between anterior and posterior compartments. **(E)** Quantification of *ex-lacZ* expression levels. Shown are the posterior/anterior (P/A) *ex-lacZ* ratios for the different genotypes analysed. Data are shown as average ± standard deviation, with all data points represented. n>9 for all genotypes. Significance was assessed using a one-way ANOVA comparing all genotypes to the *UAS-GFP* control, with Dunnett’s multiple comparisons test. *, p<0.05; ***, p<0.001; ns, non-significant. Scale bar represents 50 μm. **(F-H)** Effect of Faf catalytic activity on regulation of D localisation. XY sections of third instar wing imaginal discs, showing direct fluorescence from GFP in the control anterior compartment (F-H) or experimental posterior compartment (F’’-H’’), or Arm staining in the anterior (F’-H’) or posterior (F’’’-H’’’) compartments. Over-expression of *faf^WT^* (G’’), but not of *faf^CD^* (H’’) results in mislocalisation of D, which appears more cytoplasmic (G’’), compared to controls (F’’). No overt effects were observed for Arm in all genotypes tested (F’’’-H’’’). Scale bar represents 10 μm. **(I-L)** Faf-mediated suppression of *ft^RNAi^* growth phenotypes is dependent on its catalytic activity. Shown are adult wings from flies raised at 25°C expressing the indicated transgenes in the wing pouch under the control of *nub-Gal4*. Compared to control wings expressing *ft^RNAi^* (I), expression of *faf* suppressed the overgrowth phenotype of *ft^RNAi^* wings (J,K). This effect is dependent on the DUB activity of Faf as a catalytic mutant version of Faf (*faf^CD^*) was unable to modify the overgrowth phenotype (L). **(M)** Quantification of effect of Faf on *ft^RNAi^*-induced tissue growth. Shown are the relative adult wing sizes from flies expressing the indicated transgenes under the control of *nub-Gal4*. Data are represented as % of the average wing area of the respective controls (*nub>GFP*, average set to 100%). Data are shown as average ± standard deviation, with all data points depicted. n>19 for all genotypes. Significance was assessed using a one-way ANOVA analysis comparing all genotypes to their respective controls (*nub>GFP* or *nub>UAS-ft^RNAi^*; black or blue asterisks, respectively) with Dunnett’s multiple comparisons test. **, p<0.01; ***, p<0.001; ns, non-significant. Scale bar represents 500 μm.

### The effects of Faf are dependent on its catalytic activity

Faf is part of the DUB family of proteins, enzymes that regulate ubiquitylation levels and counteract the action of ubiquitin E3 ligases in cells ^**46**^. Given that Faf is predicted to be an active DUB, we next tested whether the role of Faf in tissue growth and modulation of Ft and Hippo signalling events was dependent on its catalytic activity. To this end, we used previously generated *UAS*-regulated transgenes encoding either WT or a catalytically inactive form of Faf (respectively, Faf^WT^ and Faf^CD^) ^**47**^ inserted at the same genomic location. First, we assessed the role of the catalytic activity of Faf in the regulation of Yki-dependent transcription using *ex-lacZ* **(Figure 7A-E)** and *HRE-DIAP1::GFP* **(Figure S6A-E)**. Over-expression of Faf resulted in a reduction in *ex-lacZ* levels when compared to the *UAS-GFP* control **(Figure 7A-C and 7E)**. Similar results were obtained when the *DIAP1::GFP* reporter was analysed **(Figure S6A-C and S6E)**. These results confirm the effect of Faf on tissue growth and support the hypothesis that, at least in part, the role of Faf involves the regulation of Hippo signalling activity. Interestingly, when we tested the catalytic mutant version of Faf (*faf^CD^*) in the same conditions, there was no effect on either the *ex-lacZ* **(Figure 7D and 7E)** or the *DIAP1::GFP* reporters **(Figure S6D and S6E)**. This strongly suggests that the role of Faf in the regulation of tissue growth and Hippo signalling is dependent on its catalytic activity and DUB function, as opposed to a potential role as a scaffolding protein bridging protein-protein interactions.

To further validate these observations, we assessed other Ft-related phenotypes, such as the regulation of D subcellular localisation. As shown in **Figures 5** and **S5**, Faf over-expression resulted in loss of D membrane polarisation (and, in some cases, loss of membrane localisation) and increased D cytoplasmic levels. These results were recapitulated when we overexpressed *faf^WT^*in the developing wing disc **(Figure 7G)**. Compared to the respective anterior compartment and the controls (*lacZ^RNAi^*), tissues where *faf^WT^* was overexpressed had D subcellular localisation defects. In contrast, when we overexpressed the *faf^CD^* mutant, we observed no overt effects on the levels or subcellular localisation of D **(Figure 7H)**. Again, this suggests that the effect of Faf on Ft signalling events is dependent on its catalytic activity.

We also directly assessed the role of Faf DUB activity in the regulation of tissue growth and on the genetic interactions with Ft. For this, we expressed Faf, Faf^WT^ and Faf^CD^ in the wing pouch using *nub-Gal4* and measured wing size as a proxy for the effects of Faf on tissue growth. When we tested the transgenes in isolation, as expected, we observed phenotypes consistent with the proposed role of Faf in the regulation of tissue growth **(Figure S6F-I)**. Adult wings from animals expressing *UAS-faf* **(Figure S6G)** or *UAS-faf^WT^* **(Figure S6H)** were significantly smaller than those of the controls (*nub>GFP*, **Figure S6F** and **Figure S6J**). Interestingly, the phenotype elicited by *UAS-faf^WT^* was more severe than *UAS-faf*, which can be explained by the fact that the transgenes have different genomic locations. Importantly, when *UAS-faf^CD^* was expressed, we did not observe tissue growth restriction but, instead, the adult wings were slightly larger than controls **(Figure S6I)**. This suggests that the catalytic activity of Faf is required for its effect on tissue growth. Moreover, the mild overgrowth of Faf^CD^ wings raises the possibility that this catalytically inactive allele may be acting as a mild dominant negative version of Faf.

Next, we assessed genetic interactions between Faf and Ft by expressing the Faf transgenes in the presence of Ft RNAi-mediated depletion **(Figure 7I-M)** or Ft over-expression **(Figure S6K-O)**. As previously shown, depletion of *ft* resulted in overgrowth phenotypes **(Figure 7I)**. In agreement with our previous results, expression of *UAS-faf* in these conditions abrogated this overgrowth phenotype **(Figure 7J)**. Similarly, expression of *UAS-faf^WT^*blocked the overgrowth caused by depletion of *ft* and, in fact, caused a significant undergrowth phenotype **(Figure 7K)**. Again, in agreement with our hypothesis, expression of the inactive form of Faf (*UAS-faf^CD^*) did not modify the phenotype of *ft^RNAi^* flies and the adult wing sizes were indistinguishable from those of controls (*nub>ft^RNAi^, GFP*) **(Figure 7L and 7M)**. We validated these observations by assessing the effect of co-expression of Ft and Faf **(Figure S6K-O)**. Ft over-expression caused a significant undergrowth phenotype when compared to controls **(Figure S6K and S6F)** and this was enhanced by co-expression of either *UAS-faf* **(Figure S6L)** or *UAS-faf^WT^* **(Figure S6M)**. In contrast, expression of the mutant *UAS-faf^CD^*resulted in a mild rescue of the *UAS-ft* adult wing phenotype **(Figure S6N and S6O)**. Together, our *in vivo* data support our conclusion that the catalytic activity of Faf is essential for its function in the regulation of Hippo signalling and tissue growth.

### Faf-mediated regulation of Ft protein is dependent on its DUB activity

Our *in vivo* results strongly suggest that Faf DUB activity is required for its function in tissue growth. To test if this effect was directly connected to Ft regulation, we assessed the effect of WT and mutant Faf on the protein levels of Ft, *in vivo* and *in vitro*. Firstly, we used *hh-Gal4* to express the *faf* transgenes in the posterior compartment of the developing wing imaginal disc and monitored Ft protein levels using a Ft-specific antibody **(Figure 8A-D)**. We observed that, in agreement with our previous observations, expression of *faf^WT^* resulted in an increase in Ft protein levels (expressed as the ratio between the levels of Ft in the posterior and anterior compartments or normalised to the corresponding Arm levels) **(Figure 8B, 8D and 8F)**, whereas the expression of the *faf^CD^* catalytic mutant had no obvious effect on Ft antibody staining **(Figure 8C, 8D and 8F)**. Importantly, despite the fact that *faf^WT^* caused significant changes in wing disc morphology, none of the Faf transgenes affected the protein levels of Arm, which was used as a control for a global effect on apical membrane proteins **(Figure 8E)**.

**Figure 8.**
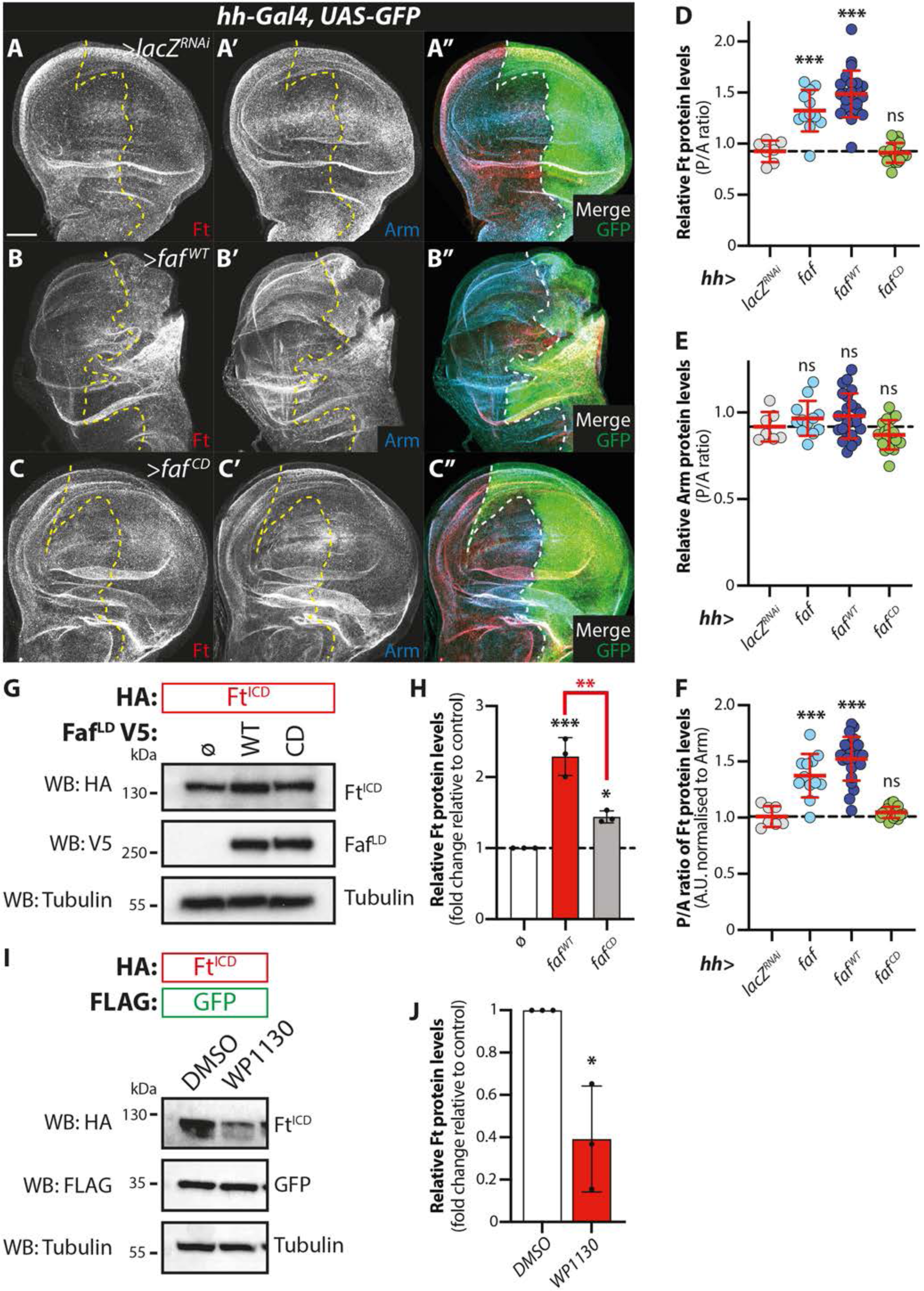
Faf regulation of Ft protein levels is DUB-dependent. **(A-C)** Faf regulates Ft protein levels *in vivo* in a DUB-dependent manner. Shown are XY confocal micrographs of third instar wing imaginal discs expressing the indicated constructs under the control of *hh-Gal4*, showing Ft antibody staining (A-C and red in A’’-C’’), Arm antibody staining (A’-C’ and cyan in A’’-C’’), and direct fluorescence from GFP (green in A’’-C’’ merged images). Compared to the controls (A, *lacZ^RNAi^*), over-expression of WT *faf* (B), but not of catalytically inactive *faf* (*faf^CD^*, C), resulted in an increase in Ft protein levels. Ventral is up in XY sections, whilst GFP marks the *hh-Gal4*-expressing posterior compartment (right). Dashed white line depicts boundary between anterior and posterior compartments. Scale bar represents 50 μm. **(D-F)** Quantification of *in vivo* Arm and Ft protein levels. Shown are the posterior/anterior (P/A) ratios for Fat protein levels (D), Arm protein levels (E), as well as the normalised Ft protein levels (F, normalised to the respective Arm levels). Data are shown as average ± standard deviation, with all data points represented. n=7, 12, 23 and 18 for *lacZ^RNAi^*, *faf*, *faf^WT^*and *faf^CD^*, respectively. Significance was assessed using a one-way ANOVA comparing all genotypes to the respective control (*hh>lacZ^RNAi^*), with Dunnett’s multiple comparisons test. **, p<0.01; ***, p<0.001. **(G-J)** Regulation of Ft protein levels in *Drosophila* S2 cells is mediated by the DUB activity of Faf. (G) HA-tagged Ft^ICD^ was co-expressed with and V5-tagged Faf (WT or a catalytically inactive version, CD) in *Drosophila* S2 cells. Cells were lysed and lysates were analysed by immunoblot using the indicated antibodies for detection of protein expression. Tubulin was used as loading control. (H) Quantification of effect of Faf catalytic activity Ft protein levels in S2 cells. Shown are the relative Ft protein levels (fold change relative to controls, which were set to 1 in cells transfected with empty plasmid (ø); Ft protein levels were normalised to their respective Tubulin control) quantified from Western blot experiments where Faf was over-expressed. Data are represented as average ± standard deviation, with all data points represented. n=3 independent experiments. Significance was assessed by a one-way ANOVA comparing all samples to the respective control (ø), with Tukey’s multiple comparisons test or an unpaired t-test. *, p<0.05 **, p<0.01; ***, p<0.001. (I) Inhibition of DUB activity affects Ft protein stability. S2 cells were treated with 5 μM of WP1130 for 6 h before cell lysis. Protein levels were analysed by Western blotting using the indicated antibodies after cell lysis. FLAG-tagged GFP and Tubulin (Tub) were used as transfection and loading controls, respectively. (J) Quantification of effect of DUB inhibition on Ft protein levels in S2 cells. Significance was assessed by an unpaired t-test. *, p<0.05.

Next, we sought to confirm these observations *in vitro* in *Drosophila* S2 cells. For that, we generated a catalytically inactive version of Faf (Faf^CD^) in our Faf^LD^ cDNA clone and tested whether its expression modulated Ft protein levels. In contrast to the WT version of Faf, which promoted the stabilisation of Ft and resulted in higher levels of Ft protein in cell lysates, expression of Faf^CD^ in S2 cells had a minimal effect on Ft levels **(Figure 8G and 8H)**. Importantly, as in this situation we assessed the levels of epitope-tagged Ft rather than its endogenous levels, the effects of Faf are likely due to post-translational modifications and not via the regulation of endogenous *ft* gene transcription and/or translation. This is consistent with the notion that Faf regulates Ft in a DUB-dependent manner. We further tested this possibility by treating *Drosophila* S2 cells with a DUB inhibitor that affects Faf function, WP1130 ^**48, 49**^ **(Figure 8I and 8J)**. WP1130-treated cells exhibited lower levels of Ft protein than cells treated with vehicle **(Figure 8I and 8J)**, reinforcing the notion that the catalytic activity of Faf is important for its modulation of Ft protein levels and, subsequently, of Ft-mediated signalling.

### Evolutionary conservation of Faf function in tissue growth regulation

Given that Faf is part of an evolutionarily conserved protein family with recognisable orthologues in other species, we next sought to determine whether the function of Faf in the regulation of Ft-mediated signalling is conserved. For this, we initially tested the effect of the mammalian orthologue of Faf, USP9X, *in vivo* in *Drosophila* tissues. Using the *en-Gal4* driver, we expressed *UAS-USP9X* in the posterior compartment of the developing wing and assessed whether this impacted on the expression of the reporters of Yki activity, *ex-lacZ* or *HRE-DIAP1::GFP* **(Figure 9A-D)**. In a similar manner to *UAS-faf^WT^*, expression of its mammalian paralog *USP9X* resulted in a reduction in the posterior/anterior (P/A) ratio of *ex-lacZ* and *DIAP1::GFP* expression **(Figure 9A-D)**. We also determined whether USP9X disrupted the subcellular localisation and polarisation of D in the wing disc. Similar to *UAS-faf^WT^*, expression of *USP9X* resulted in a disordered D localisation in the wing epithelial cells **(Figure 9E and 9F)**. Similarly to *UAS-faf* **(Figure 5E’’)**, we observed that cells expressing *UAS-USP9X* in the posterior compartment showed poor planar cell polarisation of D and, in several instances, cells appeared to display D discontinuously at the apical membrane or to lack it altogether **(Figure 9F’)**. These data suggest that, at least when overexpressed, USP9X phenocopies Faf in *Drosophila* tissues, supporting the hypothesis that the function of Faf in tissue growth is conserved in mammalian tissues.

**Figure 9.**
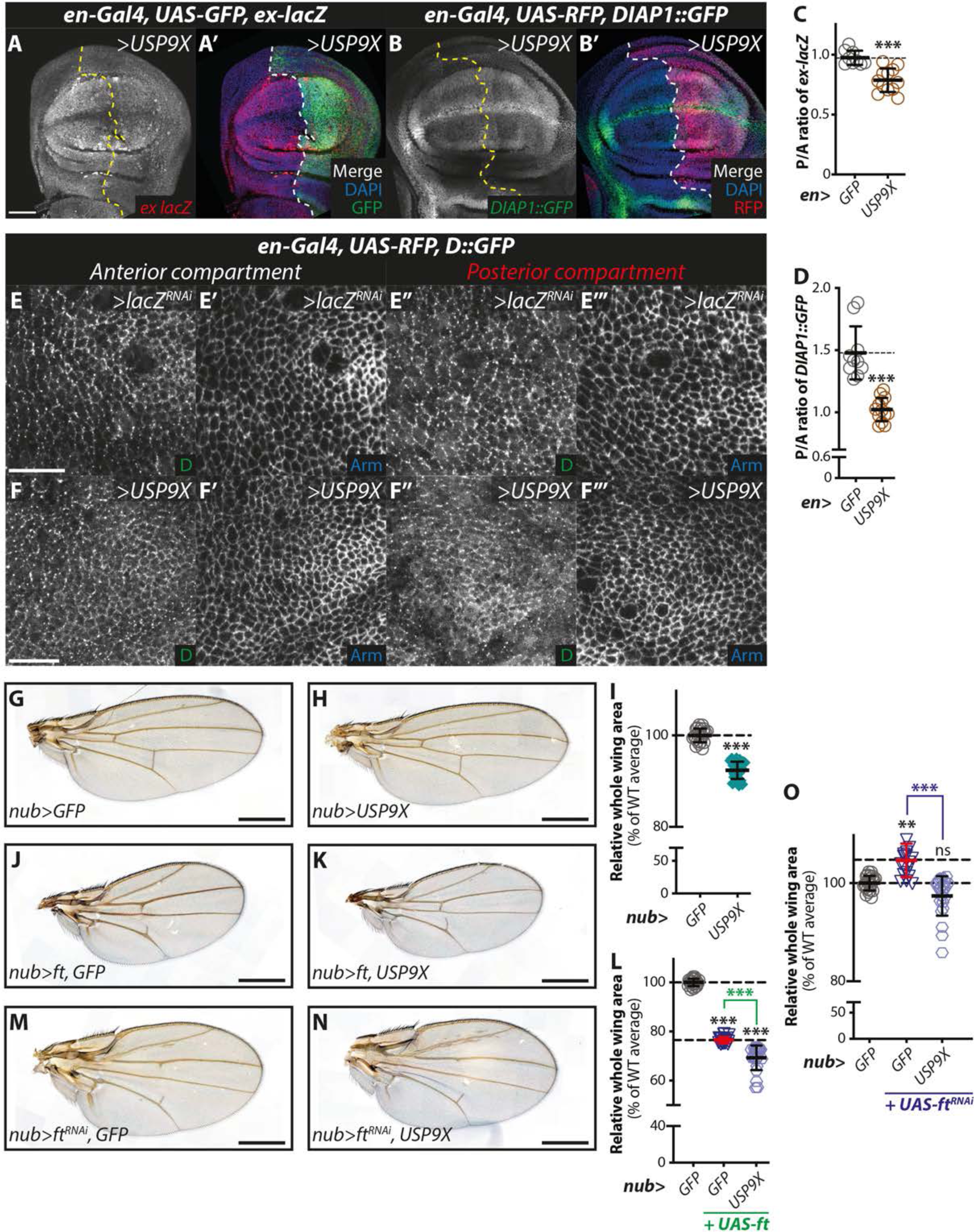
Faf-mediated regulation of Hpo and Ft signalling is conserved. **(A, B)** USP9X regulates Hpo signalling readouts *in vivo*. Shown are XY confocal micrographs of third instar wing imaginal discs carrying *ex-lacZ* (A) or *DIAP1::GFP* (B) and expressing *UAS-USP9X* under the control of *en-Gal4*, showing β-Gal antibody staining (A and red in A’ merged image) or direct fluorescence from GFP (B and green in B’ merged image). Compared to the control anterior compartment, levels of both *ex-lacZ* and *DIAP1::GFP* are reduced upon expression of *USP9X*. Ventral is up in XY sections. GFP and RFP mark the *hh-Gal4*-expressing posterior compartment (right) in A and B, respectively. Dashed lines depict boundary between anterior and posterior compartments. Scale bar represents 50 μm. **(C, D)** Quantification of Yki-mediated transcriptional reporter expression. Shown are the posterior/anterior (P/A) ratios for *ex-lacZ* (C) and *DIAP1::GFP* levels (D). Data are shown as average ± standard deviation, with all data points represented. n>9 for all conditions. Significance was assessed using an unpaired t-test, with Welch’s correction. ***, p<0.001. **(E, F)** USP9X regulates D subcellular localisation. XY sections of third instar wing imaginal discs, showing direct fluorescence from GFP in the control anterior compartment (E, F) or experimental posterior compartment (E’’, F’’), or Arm staining in the anterior (E’, F’) or posterior (E’’’, F’’’) compartments. Over-expression of *USP9X* results in mislocalisation of D (F’’), which appears less polarised at the membrane when compared to *lacZ^RNAi^* controls (E’’). No overt effects were observed for Arm (E’’’, F’’’). Scale bar represents 10 μm. **(G-O)** USP9X regulates tissue growth and modulates Ft-mediated growth phenotypes. Shown are adult wings from flies raised at 25°C expressing the indicated transgenes in the wing pouch under the control of *nub-Gal4* (*nub>*). Compared to control adult wings expressing GFP (G), USP9X-expressing wings were smaller (H). Ft over-expression reduced tissue size (J) and this was enhanced by co-expression of USP9X (K). *ft* depletion caused increased growth (M), which was abrogated by expression of USP9X (N). Quantification of the effects of USP9X on growth is shown in (I, L and O). Data are represented as % of the average wing area of the respective controls (*nub>GFP*, average set to 100%). Data are shown as average ± standard deviation, with all data points depicted. n>19 for all genotypes. Significance was assessed using one-way ANOVA analysis comparing all genotypes to the respective control (*nub>GFP* or nub>GFP, *ft^RNAi^*; black or blue asterisks, respectively), with Dunnett’s multiple comparisons test. For pairwise comparisons, unpaired t-tests with Welch’s correction were used. **, p<0.01; ***, p<0.001; ns, non-significant. Scale bar represents 500 μm.

To test this idea further, we analysed tissue growth parameters in *Drosophila* adult wings using *nub-Gal4* to express *UAS-USP9X*. In agreement with the effect of USP9X on Yki-mediated gene expression, expression of *UAS-USP9X* in the wing pouch resulted in an undergrowth phenotype **(Figure 9H)**, when compared with controls **(Figure 9G)**. Tissue growth was significantly reduced as evidenced in **Figure 9I**. We also tested whether USP9X was able to genetically interact with Ft by combining *UAS-USP9X* with Ft over-expression or RNAi-mediated depletion. Co-expression of *ft* and *USP9X* enhanced the undergrowth phenotype of *UAS-ft* wings **(Figure 9J-L)**, indicating that Ft downstream signalling is potentially more active in the presence of ectopic USP9X. In contrast to *ft* over-expression, depletion of *ft* (*UAS-ft^RNAi^*) resulted in enhanced tissue growth in the wing **(Figure 9M)**. Expression of *USP9X* in these conditions resulted in a suppression of the *UAS-ft^RNAi^* phenotype **(Figure 9N)**, and a return to WT wing tissue size **(Figure 9O)**. This is consistent with the proposed effect of Faf on the regulation of Ft protein levels and suggests that USP9X retains at least some of the functions of Faf in this context.

### USP9X-mediated regulation of Ft is conserved

Next, we assessed whether the effects seen with USP9X over-expression in *Drosophila* tissues are related to its potential regulation of Ft protein levels. To test this, we first monitored Ft protein levels in the *Drosophila* wing imaginal disc **(Figure 10A-E)**. We assessed the effect of USP9X on Ft protein levels by expressing it under the control of *hh-Gal4* and measuring Ft protein levels. Compared with the respective control (*hh>lacZRNAi*; **Figure 10A**), the levels of Ft were increased in the posterior compartment of *hh>USP9X* wing imaginal discs **(Figure 10B, 10C and 10E)**, while Arm levels were unaffected **(Figure 10A’, 10B’ and 10D)**.

**Figure 10.**
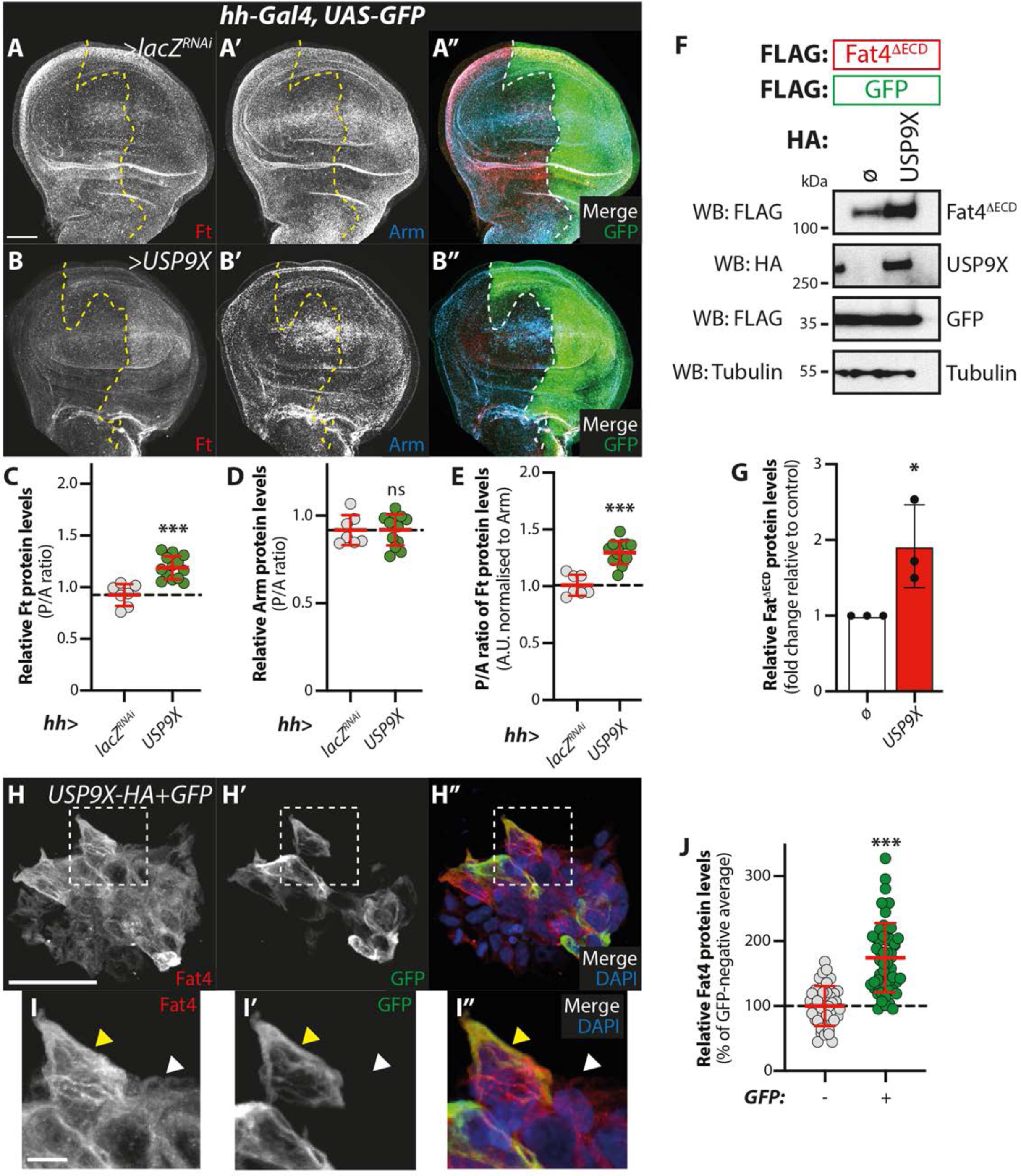
USP9X regulates Ft protein levels. **(A,B)** USP9X regulates Ft protein levels *in vivo*. Shown are XY confocal micrographs of third instar wing imaginal discs expressing the indicated constructs under the control of *hh-Gal4*, showing Ft antibody staining (A and B; and red in A’’ and B’’), Arm antibody staining (A’ and B’; and cyan in A’’ and B’’), and direct fluorescence from GFP (green in A’’ and B’’ merged images). Compared to the controls (A, *lacZ^RNAi^*), over-expression of USP9X (B) resulted in an increase in Ft protein levels. Note that panels A-A’’ are identical to Figure 8A-8A’’. Ventral is up in XY sections, whilst GFP marks the *hh-Gal4*-expressing posterior compartment (right). Dashed white line depicts boundary between anterior and posterior compartments. Scale bar represents 50 μm. **(C-E)** Quantification of *in vivo* Arm and Ft protein levels. Shown are the posterior/anterior (P/A) ratios for Fat protein levels (C), Arm protein levels (D), as well as the normalised Ft protein levels (E, normalised to the respective Arm levels). Data are shown as average ± standard deviation, with all data points represented. n=7 and 13 for *lacZ^RNAi^*and *USP9X*, respectively. Significance was assessed using an unpaired t-test, with Welch’s correction. ns, non-significant; ***, p<0.001. **(F,G)** USP9X regulates Fat4 protein levels. FLAG-tagged Fat4^ΔECD^ was co-expressed with FLAG-tagged GFP and either empty vector (ø) or HA-tagged USP9X in HEK293 cells. Cells were lysed and lysates were analysed by immunoblot using the indicated antibodies for detection of protein expression. FLAG-GFP and Tubulin were used as transfection and loading controls, respectively. (G) Quantification of effect of USP9X on Fat4^ΔECD^ protein levels in HEK293 cells. Shown are the relative Fat4 protein levels (fold change relative to controls, which were set to 1 in cells transfected with empty plasmid (ø); Fat4 protein levels were normalised to their respective Tubulin control) quantified from Western blot experiments where USP9X was over-expressed. Data are represented as average ± standard deviation, with all data points represented. n=3 independent experiments. Significance was assessed by an unpaired t-test. *, p<0.05. **(H-J)** Analysis of USP9X-dependent regulation of Fat4 protein levels via immunostaining. HEK293 cells were transfected with HA-tagged USP9X and GFP, plated in coverslips and stained with Fat4 antibody and DAPI. Shown are XY confocal images (H) and insets (I) of HEK293 cells depicting Fat4 antibody staining (grey in H and I and red in H’’ and I’’ merged images) or direct fluorescence from GFP (grey in H’ and I’ and green in H’’ and I’’ merged images). DAPI (blue in H’’ and I’’) stains nuclei. White dashed boxes in H, H’ and H’’ represent inset images I, I’ and I’’, respectively. Yellow and white arrowheads in inset images indicate GFP-positive and GFP-negative cells, respectively. Scale bar represents 50 μm in H and 10 μm in I. (J) Quantification of Fat4 protein levels in HEK293 cells transfected with USP9X. Data are represented as average ± standard deviation, with all data points represented (n=42 for GFP-negative and n=54 for GFP-positive cells). Significance was assessed by an unpaired t-test with Welch’s correction. ***, p<0.001.

We next tested whether USP9X could control Ft protein levels in mammalian cells. In mammals, there are multiple genes encoding Ft cadherins, *Fat1-4* ^**12**^. However, taking into account protein homology and function, Fat4 is the closest mammalian orthologue of Ft ^**12**^. Therefore, we assessed if the stabilisation of Ft by Faf could be recapitulated in a mammalian setting using the corresponding mammalian proteins, USP9X and Fat4. To this end, we used plasmids encoding different truncations of the Fat4 protein (Fat4^ICD^ or Fat4^ΔECD^), alongside a plasmid encoding USP9X and expressed them in mammalian HEK293 cells. Cells were transfected with Fat4 alone or in combination with USP9X and FLAG-tagged GFP was used as a transfection control. Cell lysates were analysed by Western blot analysis, which revealed that Fat4 levels were strongly increased in the presence of USP9X **(Figure 10F, 10G and S6P)**. Importantly, this effect was observed for both Fat4^ICD^ and Fat4^ΔECD^ **(Figure 10F and S6P)**. Given that both Fat4^ICD^ and Fat4^ΔECD^ largely lack the extracellular domain of Fat4, our data indicates that the effect of USP9X on the levels of Fat4 is likely to be independent of any interactions with its cognate partners that associate with the Fat4 extracellular domain. Moreover, since we assessed a FLAG-tagged Fat4 version rather than the endogenous Fat4, the USP9X-mediated regulation of Fat4 levels is predicted to be primarily the result of a post-translational mechanism, rather than an indirect effect on the expression of the *Fat4* gene.

We sought to validate our observations by directly assessing Fat4 levels in immunofluorescence experiments **(Figure 10H-J)**. For this, HEK293 cells were transfected with GFP and either Fat4 **(Figure S6Q)** or USP9X **(Figure 10H and 10I)** and stained with an anti-Fat4 antibody. Subsequently, Fat4 protein levels were analysed and compared between control GFP-negative cells and USP9X-expressing, GFP-positive cells. Analysis of our immunofluorescence experiments revealed that cells expressing GFP and USP9X exhibited higher levels of Fat4 than cells that did not express GFP **(Figure 10H-J)**. Taken together, our data indicates that, similarly to the role of Faf in *Drosophila*, USP9X stabilises Fat4 protein in mammalian cells.

## Discussion

Ft is an atypical cadherin with essential functions in tissue growth and cell polarity ^**12, 50**^. Despite intense study into its cellular role, the mechanisms regulating Ft function and its downstream effects remain relatively elusive. Ft interacts, both genetically and physically with many proteins involved in the regulation of tissue growth and planar cell polarity ^**11, 50, 51**^, but it is still unclear how these two functions and the multiple interactions are regulated and coordinated. Interestingly, several Ft-associated proteins and processes that are thought to be linked to protein ubiquitylation, such as the regulation of Ex protein stability by Dlish ^**26**^, the regulation of Wts levels and function by D ^**17, 20**^, D regulation by the F-box protein Fbxl7 ^**30, 31**^, and the regulation of D function by the E3 ligase Elgi ^**32**^. Surprisingly, despite these previous reports, the action of DUBs has not been associated with the regulation of *Drosophila* Ft. Here, we identified Fat facets (*faf*) as a novel regulator of Ft and delineated its role in the regulation of Ft protein stability and function. Our observations strongly suggest that Faf modulates the role of Ft in the regulation of tissue growth and Hippo signalling, in a manner consistent with a mechanism involving the regulation of Ft protein levels. Tissue growth phenotypes associated with increased or decreased levels of Ft in developing tissues were abrogated or enhanced when Faf levels were modified by over-expression or RNAi-mediated depletion, in agreement with a role for Faf in promoting Ft protein stability. Faf also similarly modulated the tissue growth phenotypes of Ds, the main Ft protein partner in the control of cell-cell communication and signalling, and critical for the regulation of Ft functions ^**50**^. In addition, Faf directly affected Hippo signalling readouts *in vivo*. Depletion of *faf* resulted in increased transcription of Yki target genes, indicative of defects in Hippo signalling and consistent with a reduction in Ft protein levels, as Ft is known to activate the Hippo kinase cascade, at least in part, via the regulation of Ex function ^**26, 52**^. In fact, the effect of Ft on Hippo signalling appears to be dependent on Faf function as depletion of *faf* abrogated the effect of Ft over-expression on the expression of Yki target genes. We also observed a genetic interaction between *faf* and *ft*, with loss of one allele of *faf* resulting in an enhancement of *ft* trans-heterozygous phenotypes. This further suggests that Faf function is important for the role of Ft in the regulation of tissue growth. However, we cannot exclude the hypothesis that this enhancement is due to a parallel function of Faf that impinges on imaginal disc growth.

Our results suggest that Faf mainly regulates the tissue growth function of Ft, rather than its PCP functions, since Faf affected Ft-induced tissue size but not tissue shape phenotypes. This observation was somewhat unexpected, given that Faf has been previously shown to regulate the protein levels of core PCP proteins in the wing via the regulation of Flamingo (Fmi) ^**53**^ and that the mammalian orthologue USP9X has been associated with PCP regulation via the deubiquitylation of DVL2 ^**54**^. We have not directly addressed whether Faf regulates Ft-mediated PCP downstream readouts and, therefore, cannot rule out a function for Faf in modulating Ft-related PCP functions at this stage. However, our results suggest that a putative function of Faf in regulating PCP in the context of the Ft signalling pathway is not as relevant as its involvement in tissue growth regulation. It is also possible that ubiquitylation may be a signal or a switch controlling the role of Ft and directing it toward tissue growth regulation or modulation of PCP. Nevertheless, further investigation is required to reconcile these observations.

The effects of Faf on tissue growth and regulation of Hippo signalling in *Drosophila* were recapitulated with the mammalian orthologue USP9X, suggesting that the role of Faf in regulating Ft function may be evolutionarily conserved. Indeed, USP9X has been previously associated with Hippo signalling regulation, albeit not with the regulation of the Ft orthologue Fat4. USP9X is thought to regulate Hippo signalling via the modulation of YAP1 ^**55**^, LATS ^**56, 57**^ and Angiomotin proteins ^**58, 59**^. In addition, USP9X has been shown to interact with and to deubiquitylate YAP1 ^**55**^, LATS ^**56, 57**^, WW45 ^**56**^, KIBRA ^**56**^, AMOT ^**56, 59**^ and AMOTL2 ^**58**^. It is still unclear how USP9X affects Hippo signalling, as some of its reported substrates have opposing effects on the pathway. Whilst some of the differences can be potentially explained by cell-type specific functions of USP9X, these are not sufficient to reconcile all previous observations. Moreover, none of these reports has assessed the role of USP9X in regulating Fat4 or other mammalian Ft proteins. Therefore, it remains a possibility that some of the previously reported effects of USP9X on Hippo signalling could be related to Fat4 regulation, particularly for those proteins known to be in the vicinity of Fat4 at the membrane. Interestingly, in the mouse heart, Fat4 regulates Hippo signalling and YAP1 activity via the Angiomotin protein Amotl1 ^**60**^, raising the possibility that USP9X may be required for this function. USP9X has a complex role in cancer, with both pro- and anti-tumourigenic functions and this may be related to the fact that it not only regulates Hippo signalling, but also other pathways such as TGF-β, Wnt and JAK-STAT, both in mammals ^**54, 61–65**^, as well as in *Drosophila* ^**66**^.

Our data firmly establishes Faf as a regulator of Ft function and tissue growth. However, questions remain regarding the precise molecular mechanisms involved. Our data is consistent with a direct effect of Faf on Ft, given that we observed that the two proteins can interact in co-IP experiments. In agreement with our hypothesis, S2 cell and *in vivo* data revealed that Faf controls Ft protein levels post-translationally. Moreover, the role of Faf is dependent on its catalytic activity, suggesting that Faf-mediated deubiquitylation is required for the regulation of Ft protein levels. This is further supported by the observation that blocking Faf activity using the WP1130 DUB inhibitor affects Ft protein levels. However, whilst the DUB activity of Faf seems to be required for the regulation of Ft protein levels and function, it is still unclear whether Faf directly deubiquitylates Ft or if the effect of Faf is indirect, even though the two proteins bind to each other. Analysis of a direct effect of Faf in deubiquitylating Ft is complicated by the lack of evidence regarding direct Ft ubiquitylation and the fact that no E3 ligase has been identified that targets Ft.

Previous efforts have failed to establish Elgi as a Ft E3 ligase ^**32**^ and, therefore, we have not tested whether Faf-mediated regulation of Ft is related to the E3 ligase activity of Elgi. Without a clear E3 ligase candidate to promote ectopic Ft ubiquitylation in *Drosophila* S2 cells, it would be challenging to firmly establish Faf as a direct Ft DUB. Alternative models that are not dependent on Faf directly acting on Ft are also plausible. For instance, although the precise mechanistic details remain unclear, in their role as modulators of Ft-mediated signalling, both Fbxl7 and Elgi have been proposed to regulate protein trafficking ^**30, 32**^. Therefore, it is possible that the effect of Faf is also related to protein trafficking. Indeed, it has been previously reported that the mechanism by which Faf regulates PCP in *Drosophila* is potentially related to protein trafficking, as loss of *faf* results in increased lysosomal degradation of Fmi, a possible consequence of improper vesicle accumulation of internalised Fmi ^**53**^. Moreover, Faf and its putative substrate Lqf (Liquid facets, a *Drosophila* Epsin) have been proposed to control endocytosis of several cargoes, including the Notch ligand, Delta ^**67, 68**^. Whether Faf-mediated regulation of Ft is dependent on Ft endocytosis and trafficking is still unknown, as well as the degradation mechanism, which could be mediated by the proteasome and/or lysosome.

Finally, while the level at which Faf acts within the Ft signalling pathway remains to be precisely determined, some hints come from our *in vivo* genetic interaction experiments, the analysis of D subcellular localisation, and the effects on Ft protein stability. Our data suggest that Faf likely acts at a regulatory node including Ft, Fbxl7 and App and, given the phenotypes observed, it is possible that it antagonises the action of Dlish in regulating D function. Further biochemical experiments are required to determine if the interactions between these Ft signalling components are modulated by Faf, if App-mediated palmitoylation is affected by Faf activity, or if any of these proteins is a bona fide Faf substrate.

## Materials and Methods

### Drosophila cell culture, expression constructs and chemical treatments

Work involved the use of the *Drosophila* cell line Schneider S2 (RRID:CVCL_Z232), obtained from the ATCC and screened for mycoplasma presence, showing no contamination. *Drosophila* S2 cells were grown in *Drosophila* Schneider’s medium (Thermo Fisher Scientific) supplemented with 10% (v/v) FBS, 50 μg/mL penicillin and 50 μg/mL streptomycin. Expression plasmids were transfected using Effectene transfection reagent (QIAGEN). Plasmids were generated via Gateway^®^ technology (Thermo Fisher Scientific). Open reading frames (ORFs) were PCR amplified from cDNA clones obtained from the *Drosophila* Genomics Resource Center (DGRC, https://dgrc.cgb.indiana.edu/vectors/Overview) and cloned into the pDONR207 or pDONR-Zeo Entry vectors. Destination vectors used were obtained from the *Drosophila* Gateway Vector Collection or generated in-house as previously described ^**28**^. All Entry vectors were verified by sequencing. Point mutations were generated using the Quikchange Site-Directed Mutagenesis kit (Agilent) according to the manufacturer’s instructions. The Fat^ICD^ plasmid has been previously described ^**39**^. Where indicated, inhibition of Faf was achieved by treating cells with 5 μM of the DUB inhibitor WP1130, also known as Degrasyn (Cambridge Bioscience) for 6 h before cell lysis.

### Mammalian cell culture, expression constructs and chemical treatments

Mammalian *in vitro* work involved the use of HEK293 cells (RRID:CVCL_0045), obtained from the ATCC and screened for mycoplasma presence, showing no contamination. HEK293 cells were grown in Dulbecco’s Modified Eagle Medium (DMEM, Thermo Fisher Scientific) supplemented with 10% (v/v) FBS, 50 μg/mL penicillin and 50 μg/mL streptomycin. Expression plasmids were transfected using Lipofectamine LTX Transfection Reagent (Thermo Fisher Scientific). USP9X plasmid (pCMV-HA-USP9X (DU10171)) was obtained through the MRC PPU Reagents and Services facility (MRC PPU, College of Life Sciences, University of Dundee, Scotland, mrcppureagents.dundee.ac.uk). Fat4 plasmids (pcDNA Fat4[ICD]::FLAG and pCMV5 Fat4[ΔECD]::FLAG) were a kind gift from Helen McNeill (Washington University School of Medicine, St. Louis). The pCMV-FLAG-EGFP plasmid was a kind gift from Nic Tapon (Francis Crick Institute, London).

### RNAi production and treatment

dsRNAs were synthesised using the Megascript T7 kit (Thermo Fisher Scientific) according to the manufacturer’s instructions. DNA templates for dsRNA synthesis were PCR amplified from genomic DNA or plasmids encoding the respective genes using primers containing the 5’ T7 RNA polymerase-binding site sequence. dsRNA primers were designed using the DKFZ E-RNAi design tool (https://www.dkfz.de/signaling/e-rnai3/). The following primers were used: *lacZ* (Fwd –TTGCCGGGAAGCTAGAGTAA and Rev – GCCTTCCTGTTTTTGCTCAC); *faf* (Fwd – CATCGCGTTTAGGCGAGTA and Rev – CGCACCACGCTGATGAGTA). After cell seeding, S2 cells were incubated with 20 μg dsRNA for 1 h in serum-free medium, before complete medium was added. 72 h after dsRNA treatment, cells were lysed and processed as detailed below.

### Immunoprecipitation and immunoblot analysis

For purification of FLAG-tagged proteins, cells were lysed in lysis buffer (50 mM Tris pH 7.5, 150 mM NaCl, 1% Triton X-100, 10% (v/v) glycerol, and 1 mM EDTA), to which 0.1M NaF, phosphatase inhibitors 2 and 3 (Sigma) and protease inhibitor cocktail (Complete, Roche) were added. Cell extracts were spun at 17,000 g for 10 min at 4°C. FLAG-tagged proteins were purified using anti-FLAG M2 Affinity agarose gel (Sigma) for >1h at 4°C. FLAG immunoprecipitates were then washed four times with lysis buffer before elution using 150ng/μl 3x FLAG peptide for 15-30 minutes at 4°C. Detection of purified proteins and associated complexes was performed by immunoblot analysis using chemiluminescence (Thermo Fisher Scientific). Western blots were probed with mouse anti-FLAG (M2; Sigma; RRID:AB_262044), rat anti-HA (3F10; Roche Applied Science; RRID:AB_2314622), mouse anti-V5 (Thermo Fisher Scientific; RRID:AB_2556564), or mouse anti-tubulin (E7; DSHB; RRID:AB_528499). Secondary antibodies used included HRP-conjugated sheep anti-mouse (Amersham) and HRP-conjugated goat anti-rat (Thermo Fisher Scientific). For densitometry analysis of immunoblots, X-ray blots were scanned using an Epson Perfection V700 flatbed scanner and further analysed with the Gel Analyzer function on ImageJ (RRID:SCR_003070). Alternatively, immunoblots were analysed in an ImageQuant 600 (Amersham).

### Immunostaining

Larval tissues were processed as previously described ^**69**^. Primary antibodies were incubated overnight at 4°C unless otherwise stated. Mouse anti-Armadillo (N2 7A1; DSHB; RRID:AB_528089) was used at 1:50, rat anti-Fat (kind gift from Helen McNeill) was used at 1:500 and mouse anti-β-galactosidase (Z3781, Promega; RRID:AB_430877) was used at 1:500. Anti-mouse Rhodamine Red-X-conjugated (Jackson ImmunoResearch) secondary antibodies were used at 1:500. Anti-mouse or anti-rat Alexa Fluor 647-conjugated (Jackson ImmunoResearch) secondary antibodies were used at 1:500. Secondary antibodies were incubated for at least 2 h at room temperature. After washes, tissues were stained with DAPI (1μg/mL) for 10 minutes before clearing in Vectashield (without DAPI) (H-1200, Vector Labs; RRID:AB_2336790, respectively), and mounting with Mowiol 40-88 (Sigma). Fluorescence images were acquired on Zeiss LSM710 or Zeiss LSM880 confocal laser scanning microscopes (40x or 63x objective lens).

### Mammalian cell immunofluorescence

HEK293 cells were seeded onto glass coverslips coated with Poly-L-Lysine (Sigma) and left to adhere overnight prior to transfection with plasmid DNA. 24-48 h after transfection, coverslips were washed in PBS, fixed with 4% Paraformaldehyde and permeabilised in 0.1% Triton-X-100 in PBS before incubating with the primary antibody Rabbit anti-Fat4 (PA5-116735; Thermo Fisher Scientific; RRID: AB_2901366) at a concentration of 1:100. Donkey anti-rabbit Rhodamine Red X-conjugated secondary antibody (Jackson Immunoresearch) was used at a concentration of 1:500. After washes, the cells were stained with DAPI (1μg/mL) for 5 minutes. Coverslips were then rinsed in distilled H_2_O, dried on tissue and mounted onto microslides containing a droplet of Mowiol 40-88 (Sigma). Slides were left to dry overnight before imaging on a LSM 880 confocal microscope (40x objective).

### Drosophila genetics and genotypes

Transgenic RNAi stocks were obtained from the Vienna *Drosophila* Resource Center (VDRC; RRID:SCR_013805) and the Kyoto Stock Center (DGRC; RRID:SCR_008469). Details of RNAi stocks used in the *in vivo* RNAi screens are detailed in **Table S1**. *ft^G-rv^* and *ft^8^* were obtained from Yanlan Mao (UCL). *en-Gal4, UAS-GFP, ex-lacZ* (*ex^6^*^97^) was obtained fron Nic Tapon (Francis Crick Institute). *en-Gal4, HRE-diap1::GFP* was obtained from Jin Jiang (UT Southwestern). *Fbxl7* stocks were obtained from Barry Thompson (ANU, Canberra). *en-Gal4, UAS-RFP* was obtained from Jean-Paul Vincent (Francis Crick Institute). The *faf* fly stocks *faf^B3^* (BL25100), *faf^BX3^*(BL25101), *faf^BX4^* (BL25107), *faf^FO8^* (BL25108), and *UAS-faf* (BL25102) were obtained from Bloomington. *UAS-faf^WT^*, *UAS-faf^C1677S^* (*UAS-faf^CD^*) and *UAS-USP9X* were a kind gift from Bassem A. Hassan (Paris Brain Institute, France) and have been previously described ^**47**^.

All crosses were raised at 25°C unless otherwise stated. Genotypes were as follows:

**Figure 1A,** 3A, 9G, S1D, S2A, S3A, S6F: *nub-Gal4, UAS-GFP / UAS-GFP*

**Figure 1B, S1E, S2H:** nub-Gal4, UAS-GFP / +; + / UAS-faf^RNAi^ ^55GD^ (VDRC 2955GD)

**Figure 1C, S1H, S6G:** UAS-faf / +; nub-Gal4, UAS-GFP

**Figure 1D, 9J, S6K:** nub-Gal4, UAS-ft^HA^ / UAS-GFP

**Figure 1E:** nub-Gal4, UAS-ft^HA^ / +; + / UAS-faf^RNAi^ ^55GD^ (VDRC 2955GD)

**Figure 1F, S6L:** UAS-faf / +; nub-Gal4, UAS-ft^HA^ / +

**Figure 1G, 7I, 9M:** *nub-Gal4, UAS-ft^RNAi^* (VDRC 108863KK) */ UAS-GFP*

**Figure 1H:** nub-Gal4, UAS-ft^RNAi^ (VDRC 108863KK) */ +; + / UAS-faf^RNAi^ ^55GD^* (VDRC 2955GD)

**Figure 1I, 7J:** UAS-faf / +; nub-Gal4, UAS-ft^RNAi^ (VDRC 108863KK) */ +*

**Figure 2B:** *w^iso^*

**Figure 2C:** ft^G-rv^ / ft^8^

**Figure 2D:** ft^G-rv^ / ft^8^; + / faf ^FO8^ (BL25108)

**Figure 2E:** ft^G-rv^ / ft^8^; + / faf ^BX4^ (BL25107)

**Figure 2F:** ft^G-rv^ / ft^8^; + / faf ^B3^ (BL25100)

**Figure 2G:** ft^G-rv^ / ft^8^; + / faf ^BX3^ (BL25101)

**Figure 3B:** nub-Gal4, UAS-ds / UAS-GFP

**Figure 3C:** nub-Gal4, UAS-ds / UAS-faf^RNAi^ ^79GD^ (VRDC 30679GD)

**Figure 3D:** nub-Gal4, UAS-ds / UAS-faf^RNAi^ ^KK^ (VRDC 107716KK)

**Figure 3E:** nub-Gal4, UAS-ds / +; + / UAS-faf^RNAi^ ^55GD^ (VDRC 2955GD)

**Figure 3F:** UAS-faf / +; nub-Gal4, UAS-ds / +

**Figure 3G:** nub-Gal4, UAS-Dlish^RNAi^ (VDRC 104282KK) */ UAS-GFP*

**Figure 3H:** nub-Gal4, UAS-Dlish^RNAi^ (VDRC 104282KK) */ +; + / UAS-faf^RNAi^ ^55GD^* (VDRC 2955GD)

**Figure 3I:** *UAS-faf* / +; *nub-Gal4, UAS-Dlish^RNAi^* (VDRC 104282KK) / *+*

**Figure 3J:** nub-Gal4, UAS-Dlish^RNAi^ (VDRC 104282KK) / *UAS-ft^HA^*

**Figure 3K:** nub-Gal4, UAS-Dlish^RNAi^ (VDRC 104282KK) / *UAS-ft^HA^; + / UAS-faf^RNAi^ ^55GD^* (VDRC 2955GD)

**Figure 3L:** nub-Gal4 / UAS-GFP; UAS-Fbxl7^GFP^ / +

**Figure 3M:** nub-Gal4 / +; UAS-Fbxl7^GFP^ / UAS-faf^RNAi^ ^55GD^ (VDRC 2955GD)

**Figure 3N:** nub-Gal4 / UAS-ft^HA^; UAS-Fbxl7^GFP^ / +

**Figure 3O:** nub-Gal4 / UAS-ft^HA^; UAS-Fbxl7^GFP^ / UAS-faf^RNAi^ ^55GD^ (VDRC 2955GD)

**Figure 4E, 8A, 10A:** + / UAS-lacZ^RNAi^; hh-Gal4, UAS-GFP / +

**Figure 4F:** hh-Gal4, UAS-GFP / UAS-faf^RNAi^ ^55GD^ (VDRC 2955GD)

**Figure 4G:** UAS-faf / +;; hh-Gal4, UAS-GFP / +

**Figure 4H:** UAS-ft^RNAi^ (VDRC 108863KK) */ +; hh-Gal4, UAS-GFP / +*

**Figure 4I:** UAS-ft^HA^ / +; hh-Gal4, UAS-GFP / +

**Figure 4J:** UAS-ft^HA^ / +; hh-Gal4, UAS-GFP / UAS-faf^RNAi^ ^55GD^ (VDRC 2955GD)

**Figure 5B, 7B, 9E, S5B:** en-Gal4, UAS-RFP / UAS-lacZ^RNAi^; Dachs::GFP / +

**Figure 5C:** en-Gal4, UAS-RFP / UAS-ft^HA^; Dachs::GFP / +

**Figure 5D:** en-Gal4, UAS-RFP / UAS-ft^HA^; Dachs::GFP / UAS-faf^RNAi^ ^55GD^ (VDRC 2955GD)

**Figure 5E:** UAS-faf / +; en-Gal4, UAS-RFP / +; Dachs::GFP / +

**Figure 6A, 7A:** en-Gal4, UAS-GFP, ex-lacZ (*ex^6^*^97^) */ UAS-GFP; MKRS / +*

**Figure 6B:** en-Gal4, UAS-GFP, ex-lacZ (*ex^697^*) */ UAS-hpo^RNAi^*(VDRC 104169KK); *MKRS/ +*

**Figure 6C:** en-Gal4, UAS-GFP, ex-lacZ (*ex^697^*) */ +; MKRS / UAS-faf^RNAi^ ^55GD^* (VDRC 2955GD)

**Figure 6D:** en-Gal4, UAS-GFP, ex-lacZ (*ex^697^*) */ UAS-ft^HA^; MKRS / +*

**Figure 6E:** en-Gal4, UAS-GFP, ex-lacZ (*ex^697^*) */ UAS-ft^HA^; MKRS / UAS-faf^RNAi^ ^55GD^* (VDRC 2955GD)

**Figure 6G, S6A:** en-Gal4, UAS-RFP / UAS-lacZ^RNAi^; HRE-diap1::GFP / +

**Figure 6H:** en-Gal4, UAS-RFP / UAS-hpo^RNAi^ (VDRC 104169KK)*; HRE-diap1::GFP / +*

**Figure 6I:** en-Gal4, UAS-RFP / +; HRE-diap1::GFP / UAS-faf^RNAi^ ^55GD^ (VDRC 2955GD)

**Figure 6J:** en-Gal4, UAS-RFP / UAS-ft^HA^; HRE-diap1::GFP / +

**Figure 6K:** en-Gal4, UAS-RFP / UAS-ft^HA^; HRE-diap1::GFP / UAS-faf^RNAi^ ^55GD^ (VDRC 2955GD)

**Figure 7B:** UAS-faf / +; en-Gal4, UAS-GFP, ex-lacZ (*ex^697^*) */ UAS-GFP; MKRS / +*

**Figure 7C:** en-Gal4, UAS-GFP, ex-lacZ (*ex^697^*) */ UAS-GFP; MKRS / UAS-faf^WT^*

**Figure 7D:** en-Gal4, UAS-GFP, ex-lacZ (*ex^697^*) */ UAS-GFP; MKRS / UAS-faf^CD^* (*faf^C1677S^*)

**Figure 7F:** en-Gal4, UAS-RFP / UAS-lacZ^RNAi^; Dachs::GFP / +

**Figure 7G:** en-Gal4, UAS-RFP / UAS-lacZ^RNAi^; Dachs::GFP / UAS-faf^WT^

**Figure 7H:** en-Gal4, UAS-RFP / UAS-lacZ^RNAi^; Dachs::GFP / UAS-faf^CD^ (*faf^C1677S^*)

**Figure 7K:** nub-Gal4, UAS-ft^RNAi^ (VDRC 108863KK) */ +; + / UAS-faf^WT^*

**Figure 7L:** nub-Gal4, UAS-ft^RNAi^ (VDRC 108863KK) */ +; + / UAS-faf^CD^* (*faf^C1677S^*)

**Figure 8B:** hh-Gal4, UAS-GFP / UAS-faf^WT^

**Figure 8C:** hh-Gal4, UAS-GFP / UAS-faf^CD^ (*faf^C1677S^*)

**Figure 9A:** en-Gal4, UAS-GFP, ex-lacZ (*ex^697^*) */ UAS-GFP; MKRS / UAS-USP9X*

**Figure 9B:** en-Gal4, UAS-RFP / +; HRE-diap1::GFP / UAS-USP9X

**Figure 9F:** en-Gal4, UAS-RFP / UAS-lacZ^RNAi^; Dachs::GFP / UAS-USP9X

**Figure 9H:** nub-Gal4, UAS-GFP / +; + / UAS-USP9X

**Figure 9K:** nub-Gal4, UAS-ft^HA^ / +; + / UAS-USP9X

**Figure 9N:** nub-Gal4, UAS-ft^RNAi^ (VDRC 108863KK) */ +; + / UAS-USP9X*

**Figure 10B:** hh-Gal4, UAS-GFP / UAS-USP9X

**Figure S1F:** nub-Gal4, UAS-GFP / UAS-faf^RNAi^ ^KK^ (VDRC 107716KK)

**Figure S1G:** nub-Gal4, UAS-GFP / UAS-faf^RNAi^ ^79GD^ (VDRC 30769GD)

**Figure S3B:** nub-Gal4, UAS-ex^2^ / UAS-GFP

**Figure S3C:** nub-Gal4, UAS-ex^2^ / +; + / UAS-faf^RNAi^ ^55GD^ (VDRC 2955GD)

**Figure S3D:** nub-Gal4, UAS-ex^2^ / UAS-ft^HA^

**Figure S3E:** nub-Gal4, UAS-ex^2^ / UAS-ftHA; + / UAS-faf^RNAi^ ^55GD^ (VDRC 2955GD)

**Figure S3F:** nub-Gal4, UAS-elgi^RNAi^ (VDRC 109617KK) */ UAS-GFP*

**Figure S3G:** nub-Gal4, UAS-elgi^RNAi^ (VDRC 109617KK) */ +; + / UAS-faf^RNAi^ ^55GD^* (VDRC 2955GD)

**Figure S3H:** UAS-faf / +; nub-Gal4, UAS-elgi^RNAi^ (VDRC 109617KK) */ +*

**Figure S3I:** nub-Gal4, UAS-elgi^RNAi^ (VDRC 109617KK) */ UAS-ft^HA^*

**Figure S3J:** nub-Gal4, UAS-elgi^RNAi^ (VDRC 109617KK) */ UAS-ft^HA^; + / UAS-faf^RNAi^ ^55GD^* (VDRC 2955GD)

**Figure S3K:** nub-Gal4, UAS-app^RNAi^ (VDRC 32863GD) */ UAS-GFP*

**Figure S3L:** nub-Gal4, UAS-app^RNAi^ (VDRC 32863GD) */ +; + / UAS-faf^RNAi^ ^55GD^* (VDRC 2955GD)

**Figure S3M:** UAS-faf / +; nub-Gal4, UAS-app^RNAi^ (VDRC 32863GD) */ +*

**Figure S3N:** nub-Gal4, UAS-app^RNAi^ (VDRC 32863GD) */ UAS-ft^HA^*

**Figure S3O:** nub-Gal4, UAS-app^RNAi^ (VDRC 32863GD) */ UAS-ft^HA^; + / UAS-faf^RNAi^ ^55GD^* (VDRC 2955GD)

**Figure S5C:** en-Gal4, UAS-RFP / UAS-ft^RNAi^ (VDRC 108863KK)*; Dachs::GFP / +*

**Figure S5D:** en-Gal4, UAS-RFP / +; Dachs::GFP / UAS-faf^RNAi^ ^55GD^ (VDRC 2955GD)

**Figure S6B:** UAS-faf / +; en-Gal4, UAS-RFP / +; HRE-diap1::GFP / +

**Figure S6C:** en-Gal4, UAS-RFP / +; HRE-diap1::GFP / UAS-faf^WT^

**Figure S6D:** en-Gal4, UAS-RFP / +; HRE-diap1::GFP / UAS-faf^CD^ (*faf^C1677S^*)

**Figure S6H:** nub-Gal4, UAS-GFP / +; + / UAS-faf^WT^

**Figure S6I:** nub-Gal4, UAS-GFP / +; + / UAS-faf^CD^ (*faf^C1677S^*)

**Figure S6M:** nub-Gal4, UAS-ft^HA^ / +; + / UAS-faf^WT^

**Figure S6N:** nub-Gal4, UAS-ft^HA^ / +; + / UAS-faf^CD^ (*faf^C1677S^*)

### Immunofluorescence quantification and statistical analyses

For quantification of Ft and Arm protein levels *in vivo* **(Figure 8D-F, Figure S4E-G, Figure 10C-E)**, relative *ex-lacZ* **(Figure 6F, 7E and 9C)** or *diap1::GFP* levels **(Figure 6L, 9D and S6E)**, ratios in posterior versus anterior compartment (P/A) were calculated by manually drawing around each compartment of the wing disc in maximum intensity projections, then measuring the mean grey pixel value in Fiji (RRID:SCR_002285). Significance was calculated using a Brown-Forsythe and Welch ANOVA comparing all genotypes to their respective controls (for Ft and Arm protein levels), or a one-way ANOVA comparing all means to the *en-Gal4* control (*UAS-GFP* for *ex-lacZ*; *UAS-lacZ^RNAi^* for *diap1::GFP*), both with Dunnett’s multiple comparisons test. Pairwise comparisons between other samples were performed using unpaired t-tests or unpaired t-tests with Welch’s correction.

### Analysis of genetic interactions in Drosophila adult wings

For analysis of genetic interactions in the *Drosophila* wing, flies with the genotypes of interest were collected and preserved in 70% EtOH for at least 24 h. Adult wing samples were prepared as previously described **^70^**. Briefly, wings were removed in 100% isopropanol, mounted in microscope slides using Euparal (Anglian Lepidopterist Supplies) as mounting medium and baked at 65°C for at least 5 h. Adult wing images were captured using a Pannoramic 250 Flash High Throughput Scanner (3DHISTECH) and extracted using the Pannoramic Viewer software (3DHISTECH). Wing area was quantified using Fiji (the alula and costal cell of the wing were both excluded from the analysis). Images were processed using Adobe Photoshop (RRID:SCR_014199).

### Statistical Analysis

Statistical analyses were performed using GraphPad Prism 8, 9 and 10 (RRID:SCR_002798). Significance values corresponding to comparisons between two groups were calculated using unpaired t-test or unpaired t-test with Welch’s correction. Significance values corresponding to comparisons between multiple groups were calculated using one-way ANOVA, Brown-Forsythe and Welch ANOVA or Kruskal-Wallis ANOVA analyses with Dunnett’s, Tukey’s or Dunn’s multiple comparisons tests. p-values: ns denotes not significant; * denotes p<0.05; ** denotes p<0.01; *** denotes p<0.001.

## Acknowledgements

We thank the Vienna *Drosophila* Resource Center for providing transgenic RNAi fly stocks used in this study. Stocks obtained from the Bloomington *Drosophila* Stock Center (NIH P40OD018537) were used in this study. The antibodies E7 (DSHB Hybridoma Product E7), 2A1 (DSHB Hybridoma Product 2A1) and N2 7A1 (DSHB Hybridoma Product N2 7A1 Armadillo) were deposited by M. Klymkowsky, R. Holmgren and E. Wieschaus to the Developmental Studies Hybridoma Bank, created by the NICHD of the NIH and maintained at The University of Iowa, Department of Biology. We thank N. Tapon, H. McNeill, B. Thompson, Y. Mao, J. P. Vincent and B. Hassan for providing fly stocks. We thank Linda Hammond for assistance with microscopy. We thank C. Mohanathas for assistance in preliminary experiments. We thank members of the Ribeiro lab for helpful discussions and N.Tapon, M. Holder, S.A. Martin, and H. McNeill for critical reading of the manuscript. The authors declare no conflicts of interest. This work was supported by funding from Cancer Research UK (C16420/A18066), The Academy of Medical Sciences/Wellcome Trust Springboard Award (SBF001/1018) and from the Biotechnology and Biological Sciences Research Council (BB/T004576/1). L.E.D. was supported by a Biotechnology and Biological Sciences Research Council LIDo PhD studentship (BB/M009513/1).

