## Supplemental Figure legends for "The deubiquitylating enzyme Fat Facets promotes Fat signalling and restricts tissue growth"

**Supplementary Figure 1 – *In vivo* RNAi screens to identify Ft signalling regulators**

**(A)** Schematic representation of the *in vivo* RNAi genetic screens performed. Temporal and spatial control of gene expression was achieved by using the *nub-Gal4* driver (*nub*>), such that, in the developing wing, expression of *UAS-RNAi* genes (targeting DUBs, deneddylases and deSUMOylases) is limited to the pouch region. **(B, C)** Quantification of relative adult wing sizes from *nub-Gal4* (*nub*>; B) and *nub-Gal4, UAS-ft* (*nub>ft*; C) screens. Data are represented as % of the average wing area of the respective controls. (*nub>GFP*, average set to 100%). Data are shown as average ± standard deviation, with all data points depicted. Significance was assessed using Brown-Forsythe and Welch ANOVA analyses comparing all genotypes to the respective control, with Dunnett's multiple comparisons test. *, $p<0.05$; **, $p<0.01$; ***, $p<0.001$. Dashed black line indicates 100% (average of the *nub>GFP* control), while red dashed line indicates the average size of adult wings from *nub>ft, GFP* flies. † depicts instances where fly crosses resulted in lethality. **(D-G)** Validation of *faf* as a *nub-Gal4* screen hit. Shown are adult wings from flies raised at 25ºC expressing the indicated transgenes in the wing pouch under the control of *nub-Gal4*. Compared to control adult wings expressing GFP (D), depletion of *faf* in the developing wing resulted in increased tissue growth (E, F and G). **(H)** In contrast to $faf^{RNAi}$, *faf* over-expression resulted in an undergrowth phenotype. Scale bar represents 500 μm.

**Supplementary Figure 2 – Characterisation of morphological traits in adult wings**

**(A)** Representative image of control adult wing (from *nub>GFP* female). **(B)** Schematic representation of the defining axes of the adult wing. AP denotes anterior-posterior, while PD denotes proximal-distal. **(B')** Schematic representation of measurements of wing shape, by analysis of a selected area of the adult wing (represented by the red area) or the analysis of a fitted ellipse of the wing area (represented by the green oval). **(C-E)** Quantification of wing shape. Data is represented as the ratio between the length of the PD axis and the length of the AP axis (C), or as circularity measures of the area selection (D) or of a fitted ellipse of the wing area (E). All data is represented average ± standard deviation, with all data points depicted. Vertical dashed lines separate different genotype conditions (*nub>*, *nub>ft* or *nub>ft*$^{RNAi}$). Significance was assessed using Kruskal-Wallis ANOVA analyses comparing all genotypes to the respective control (*nub>GFP, nub>ft* or *nub>ft*$^{RNAi}$; black, blue or red asterisks, respectively), with Dunn's multiple comparisons test. *, $p<0.05$; **, $p<0.01$; ***, $p<0.001$. ns, non-significant. $n>19$ for all genotypes. **(F-I)** Scoring of morphological defects in wing vein pattern. (F) Schematic representation of control adult wing, with the normal pattern of venation. (G) Schematic representation of adult wing with a defect in L2. (H) Representative example of an adult wing with L2 defect (from *nub-Gal4, UAS-faf*$^{RNAi\ 55GD}$ female). **(I)** Table summarising the frequency of L2 defects in selected genotypes. Shown are the frequencies of adult wings with (red) or without a defect in the L2 vein (grey) in the different genotypes indicated. Scale bar represents 500 $\mu$m.

**Supplementary Figure 3 – Genetic interactions between *faf* and Hpo and Ft pathway genes**

**(A-O)** Genetic interactions between *faf* and *ex* (A-E), *elgi* (F-J) or *app* (K-O). Shown are adult wings from flies raised at 25ºC expressing the indicated transgenes in the wing pouch under the control of *nub-Gal4* (*nub*>). Over-expression of $ex^2$ (mild *ex* allele) under the control of *UAS* sequences resulted in a dramatic decrease in wing size (B) compared to WT wings (A), which was partially suppressed by *faf* depletion (C). In contrast, expression of *ft* enhanced the $ex^2$ undergrowth phenotype (D), and this was counteracted by depletion of *faf* (E). Depletion of *elgi* ($elgi^{RNAi}$) resulted in a mild decrease in tissue growth (F), which was abrogated when *faf* was co-depleted (G). Over-expression of either *faf* (H) or *ft* (I) enhanced the undergrowth phenotype of $elgi^{RNAi}$-expressing flies and the *ft* size phenotype was rescued by $faf^{RNAi}$ (J). Depletion of *app* ($app^{RNAi}$) resulted in a mild decrease in tissue growth (K), which was not affected by modulation of *faf* levels (L,M). Expression of *ft* enhanced the undergrowth phenotype (N) and this was rescued by depletion of *faf* (O). **(P-R)** Quantification of relative adult wing sizes. Data are represented as % of the average wing area of control wings (*nub>GFP*, which were set as 100%.). Data are shown as average ± standard deviation, with all data points represented. n>19 for all genotypes. Black dashed lines represent average size of controls (100%), whilst red dashed lines indicate average size of adult wings from flies expressing $ex^2$, $elgi^{RNAi}$ or $app^{RNAi}$ under the control of *nub-Gal4*. Significance was assessed using a one-way ANOVA comparing all genotypes to their respective controls (*nub>GFP*, *nub>UAS-*$ex^2$, *nub>UAS-*$elgi^{RNAi}$, *nub>UAS-*$app^{RNAi}$, black, orange, brown and blue asterisks, respectively), with Dunnett's multiple comparisons test. *, p<0.05; **, p<0.01; ***, p<0.001; ns, non-significant. Scale bar represents 500 μm.

**Supplementary Figure 4 – Faf-mediated regulation of Ft protein levels**

**(A)** Mapping interaction between Faf and Ft. HA-tagged $Ft^{ICD}$ and GFP were co-expressed with FLAG-tagged GFP, $Faf^{LD}$ or Faf truncations in *Drosophila* S2 cells. Cells were lysed and lysates were subjected to co-immunoprecipitation using FLAG agarose beads. Lysates were analysed by immunoblot using the indicated antibodies for detection of protein expression and co-purification. HA-tagged GFP and Tubulin (Tub) were used as transfection and loading controls, respectively. **(B,C)** Quantification of effect of Faf modulation in S2 cells on Ft protein levels. Shown are the relative Ft protein levels (fold change relative to controls, which were set to 1; Ft protein levels were normalised to their respective Tubulin control) quantified from Western blot experiments where Faf was overexpressed (B) or depleted (C). Data are represented as average ± standard deviation, with all data points represented. n=3 independent experiments. Significance was assessed by unpaired t-test with Welch's correction. *, $p<0.05$ **(D)** Validation of *faf* RNAi efficiency. S2 cells were treated with dsRNAs 24 h before co-transfection with the indicated constructs. RNAi specificity and efficiency were analysed by Western blotting using the indicated antibodies. Note that *faf* dsRNA targeting sequences overlap with $Faf^{LD}$ and $Faf^{C\text{-}ter}$, but not with the $Faf^{N\text{-}ter}$ construct and, as expected, only the latter was not affected by the dsRNA. FLAG-tagged GFP and Tubulin (Tub) were used as transfection and loading controls, respectively. **(E-G)** Quantification of *in vivo* Arm and Ft protein levels in the genotypes from Figure 4. Shown are the posterior/anterior (P/A) ratios for Arm protein levels (E), Fat protein levels (F), as well as the normalised Ft protein levels (G, normalised to the respective Arm levels). Data are shown as average ± standard deviation, with all data points represented. n>8 for all genotypes. Significance was assessed using a Brown-Forsythe and Welch ANOVA comparing all genotypes to their respective controls, with Dunnett's multiple comparisons test. **, $p<0.01$; ***, $p<0.001$.

**Supplementary Figure 5 – *In vivo* regulation of D subcellular localisation**

**(A)** Schematic representation of subcellular localisation of Ft signalling components and experimental setting. Note this is identical to Figure 5A. **(B-D)** Effect of *ft* and *faf* depletion on D subcellular localisation. XY sections of third instar wing imaginal discs, showing direct fluorescence from GFP in the control anterior compartment (B-D) or experimental posterior compartment (B''-D''), or Arm staining in the anterior (B'-D') or posterior (B'''-D''') compartments. Cartoons represent expected results for each genotype tested. Depletion of *ft* or *faf* results in mislocalisation of D, which appears less polarised at the membrane (C'' and D''). No overt effects were observed for Arm in all genotypes tested (B'''-E'''). Scale bar represents 10 μm.

**Supplementary Figure 6 – The catalytic activity of Faf is required for its function in the regulation of Hpo signalling and tissue growth**

**(A-D)** Faf-mediated regulation of Yki target gene expression is dependent on its catalytic activity. XY confocal sections of third instar wing imaginal discs carrying *DIAP1::GFP* (A-D, shown in green in A'-D' merged images), in which *en-Gal4* was used to drive expression of *UAS-lacZ*$^{RNAi}$ (A), *UAS-faf* (B), *UAS-faf*$^{WT}$ (C) or *UAS-faf*$^{CD}$ (D). Compared to controls (A), *faf* over-expression (B,C) resulted in a decrease in the P/A ratio of *DIAP1::GFP*, indicating lower levels of Yki-mediated transcription. Expression of a Faf catalytic mutant had no effect (D). RFP (red in A'-D' merged images) indicates posterior compartment where transgenes are expressed. DAPI (blue) stains nuclei. Dashed lines indicate boundary between anterior and posterior compartments. **(E)** Quantification of *DIAP1::GFP* expression levels. Shown are the posterior/anterior (P/A) *DIAP1::GFP* ratios for the different genotypes analysed. Data are shown as average ± standard deviation, with all data points represented. n>8 for all genotypes. Significance was assessed using a one-way ANOVA comparing all genotypes to the *UAS-lacZ*$^{RNAi}$ control, with Dunnett's multiple comparisons test. ***, p<0.001; ns, non-significant. Scale bar represents 50 μm. **(F-I)** Effect of Faf catalytic activity on tissue growth. Shown are adult wings from flies raised at 25ºC expressing the indicated transgenes in the wing pouch under the control of *nub-Gal4*. Compared to control wings expressing GFP (F), Faf expression caused reduced growth in a DUB-dependent manner (G-I). **(J)** Quantification of effect of Faf on tissue growth. Shown are the relative adult wing sizes from flies expressing the indicated transgenes under the control of *nub-Gal4*. Data are represented as % of the average wing area of the respective controls (*nub>GFP*, average set to 100%). Data are shown as average ± standard deviation, with all data points depicted. n>18 for all genotypes. Significance was assessed using a one-way ANOVA analysis comparing all genotypes to the respective control (*nub>GFP*) with Dunnett's multiple comparisons test. *, p<0.05;

***, $p<0.001$. **(K-N)** Effect of Faf on Ft-mediated regulation of growth is DUB-dependent. Shown are adult wings from flies raised at 25ºC expressing the indicated transgenes in the wing pouch under the control of *nub-Gal4*. Compared to control wings expressing *UAS-ft* (K), Faf expression enhanced Ft-mediated growth phenotypes in a DUB-dependent manner (L-N). **(O)** Quantification of effect of Faf on Ft-mediated growth phenotypes. Shown are the relative adult wing sizes from flies expressing the indicated transgenes under the control of *nub-Gal4*. Data are represented as % of the average wing area of the respective controls (*nub>GFP*, average set to 100%). Data are shown as average ± standard deviation, with all data points depicted. $n>19$ for all genotypes. Significance was assessed using a one-way ANOVA analysis comparing all genotypes to their respective controls (*nub>GFP* or *nub>UAS-ft*; black and green asterisks, respectively) with Dunnett's multiple comparisons test. *, $p<0.05$; ***, $p<0.001$. Scale bar represents 500 µm. **(P)** USP9X regulates Fat4$^{ICD}$ protein levels. FLAG-tagged Fat4$^{ICD}$ was co-expressed with FLAG-tagged GFP and either empty vector (ø) or HA-tagged USP9X in HEK293 cells. Cells were lysed and lysates were analysed by immunoblot using the indicated antibodies for detection of protein expression. FLAG-GFP and Tubulin were used as transfection and loading controls, respectively. **(Q)** Validation of Fat4 antibody in HEK293 cells. HEK293 cells were transfected with FLAG-tagged Fat4 and GFP, plated in coverslips and stained with Fat4 antibody and DAPI. Shown are XY confocal sections of transfected HEK293 cells depicting Fat4 antibody staining (grey in Q and red in Q'' merged image) or direct fluorescence from GFP (grey in Q' and green in Q'' merged image). (Q'') Merged image depicting Fat4 and GFP signals, as well as DAPI (blue) staining nuclei.

**Table S1 – *In vivo* RNAi screen information**

| DUB family | CG# | Gene name | RNAi line |
|---|---|---|---|
| USP | *CG12082* | *Usp5* | NIG 12082R-2 |
| USP | *CG14619* | *Usp2* | VDRC 104382 KK |
| USP | *CG15817* | *Usp1* | VDRC 41605 GD |
| USP | *CG3016* | *Usp30* | VDRC 7090 GD |
| USP | *CG30421* | *Usp15-31* | VDRC 33726 GD |
| USP | *CG32479* | *Usp10* | VDRC 37859 GD |
| USP | *CG4165* | *Usp16-45* | VDRC 110286 KK |
| USP | *CG5384* | *Usp14* | VDRC 110227 KK |
| USP | *CG5794* | *puffyeye (puf)* | VDRC 27517 GD |
| USP | *CG5798* | *Usp8* | VDRC 107623 KK |
| USP | *CG7023* | *Usp12-46* | VDRC 100586 KK |
| USP | *CG8334* | *Usp32* | VDRC 18981 GD |
| USP | *CG8494* | *Usp20-33* | VDRC 42609 GD |
| USP | *CG8830* | *DUBAI* | VDRC 28960 GD |
| USP | *CG5603* | *CYLD* | VDRC 15340 GD |
| USP | *CG2904* | *echinus (ec)* | VDRC 106671 KK |
| USP | *CG1945* | *fat facets (faf)* | VDRC 30679 GD |
| USP | *CG4166* | *non-stop (not)* | VDRC 45775 GD |
| USP | *CG5486* | *Usp47* | VDRC 26027 GD |
| USP | *CG5505* | *scrawny (scny)* | VDRC 105989 KK |
| USP | *CG1490* | *Usp7* | VDRC 110324 KK |
| USP | *CG7288* | *Usp39* | NIG 7288R-1 |
| USP | *CG8232* | *PAN2* | NIG 8232R-1 |
| UCH | *CG4265* | *Uch* | VDRC 103614 KK |
| UCH | *CG3431* | *Uch-L5* | VDRC 103481 KK |
| UCH | *CG1950* | *Uch-L5R* | NIG 1950R-1 |
| UCH | *CG8445* | *calypso (caly)* | VDRC 47743 GD |
| JAMM | *CG2224* | *CG2224* | VRDC 108622 KK |
| JAMM | *CG4751* | *CG4751* | VDRC 45530 GD |
| JAMM | *CG14884* | *CSN5* | NIG 14884-R1 |
| JAMM | *CG8877* | *Prp8* | VDRC 18565 GD |
| JAMM | *CG18174* | *Rpn11* | VDRC 19272 GD |
| JAMM | *CG4673* | *Npl4* | CG4673R-2 |
| JAMM | *CG8335* | *eIF3f2* | VDRC 108169 KK |
| JAMM | *CG9769* | *eIF3f1* | VDRC 101465 KK |
| JAMM | *CG6932* | *CSN6* | VDRC 105385 KK |
| JAMM | *CG9124* | *eIF3h* | VDRC 36087 GD |
| JAMM | *CG3416* | *Rpn8* | VDRC 26183 GD |
| OTU | *CG12743* | *otu* | VDRC 108845 KK |
| OTU | *CG7857* | *CG7857* | NIG 7857R-2 |
| OTU | *CG3251* | *CG3251* | VDRC 100532 KK |
| OTU | *CG4968* | *CG4968* | VDRC 21978 GD |
| OTU | *CG4603* | *Yod1* | VDRC 21893 GD |
| OTU | *CG6091* | *Duba* | VDRC 109912 KK |
| OTU | *CG9448* | *trabid (trbd)* | VDRC 24030 GD |
| JOSEPHIN | *CG3781* | *Josd* | VDRC 108379 KK |
| ULP | *CG12359* | *Ulp1* | VDRC 106625 KK |
| ULP | *CG12717* | *pirate (pira)* | VDRC 106239 KK |
| ULP | *CG1503* | *CG1503* | VDRC 32349 GD |
| ULP | *CG8493* | *Den1* | VDRC 100591 KK |
| ULP | *CG10107* | *veloren (velo)* | VDRC 103524 KK |
| ULP | *CG32110* | *CG32110* | VDRC 107634 KK |
| MCPIP | *CG10889* | *Regnase-1* | NIG 10889R-2 |

List of genes and RNAi lines tested in the *in vivo* RNAi screen. Shown are the DUB families, CG number, gene names and RNAi fly stock information. USP, UCH, JAMM,

OTU, JOSEPHIN, ULP and MCPIP denote ubiquitin-specific protease, ubiquitin C-terminal hydrolase, Jab1/Mov34/Mpr1 Pad1 N-terminal+ domain metalloprotease, ovarian tumour, Machado-Josephin domain, ubiquitin-like specific proteases and Monocyte Chemotactic Protein-Induced Protein, respectively.
