## Supplementary figures and images for "The deubiquitylating enzyme Fat Facets promotes Fat signalling and restricts tissue growth"

### Supplemental Figure S1

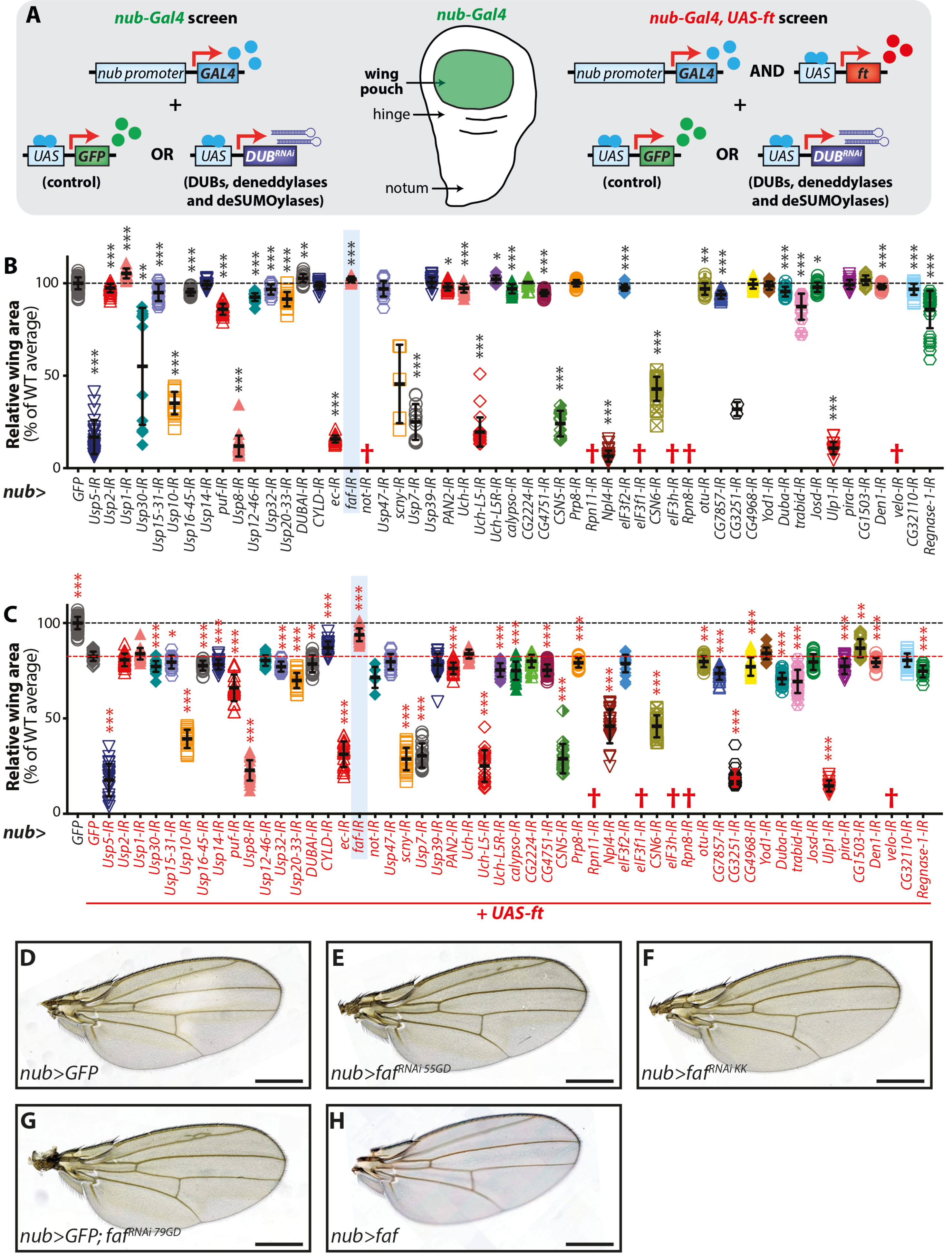


**Dawson et al. - Supplementary Figure 1**

### Supplemental Figure S2

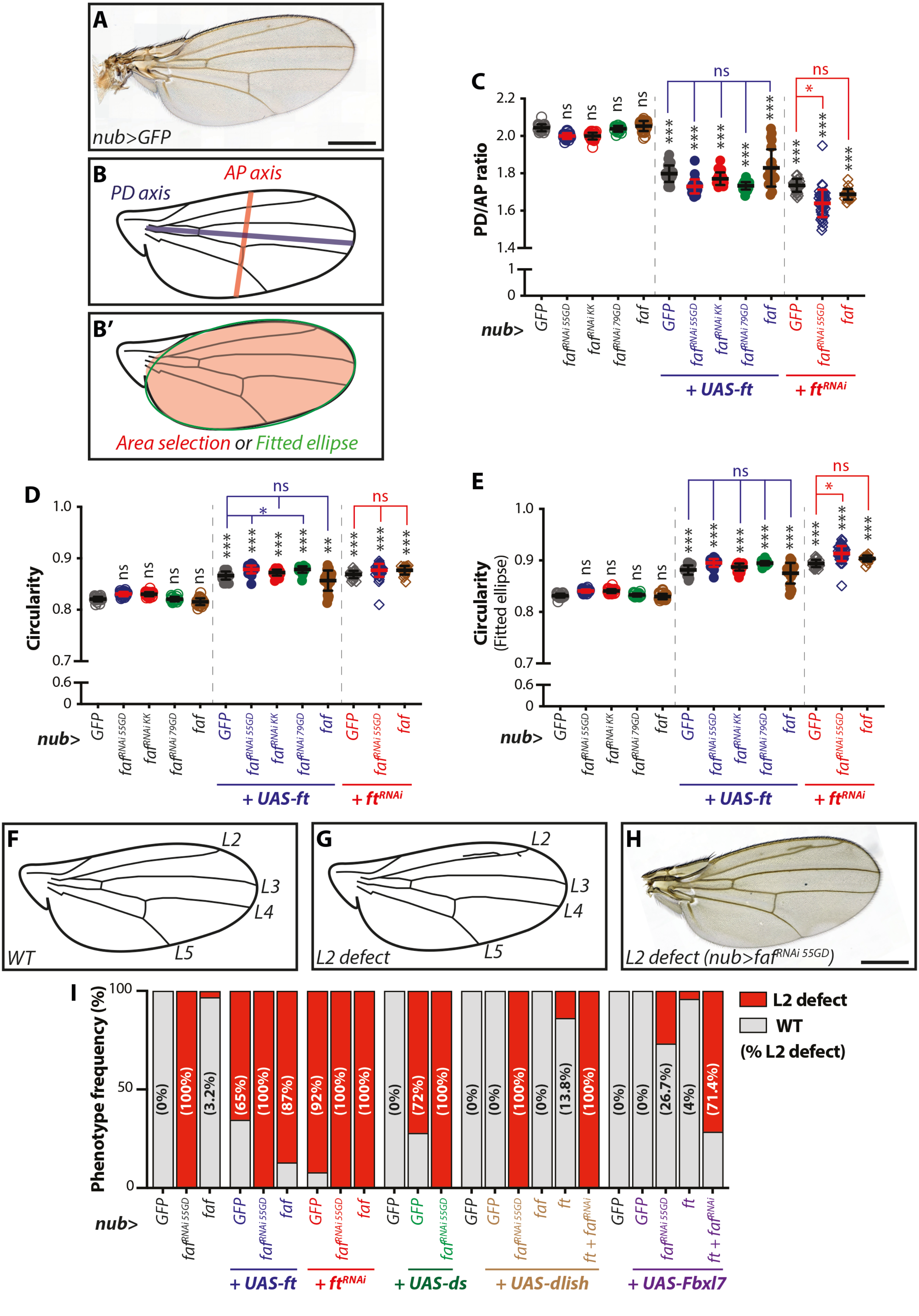


Dawson et al. - Supplementary Figure 2

### Supplemental Figure S3

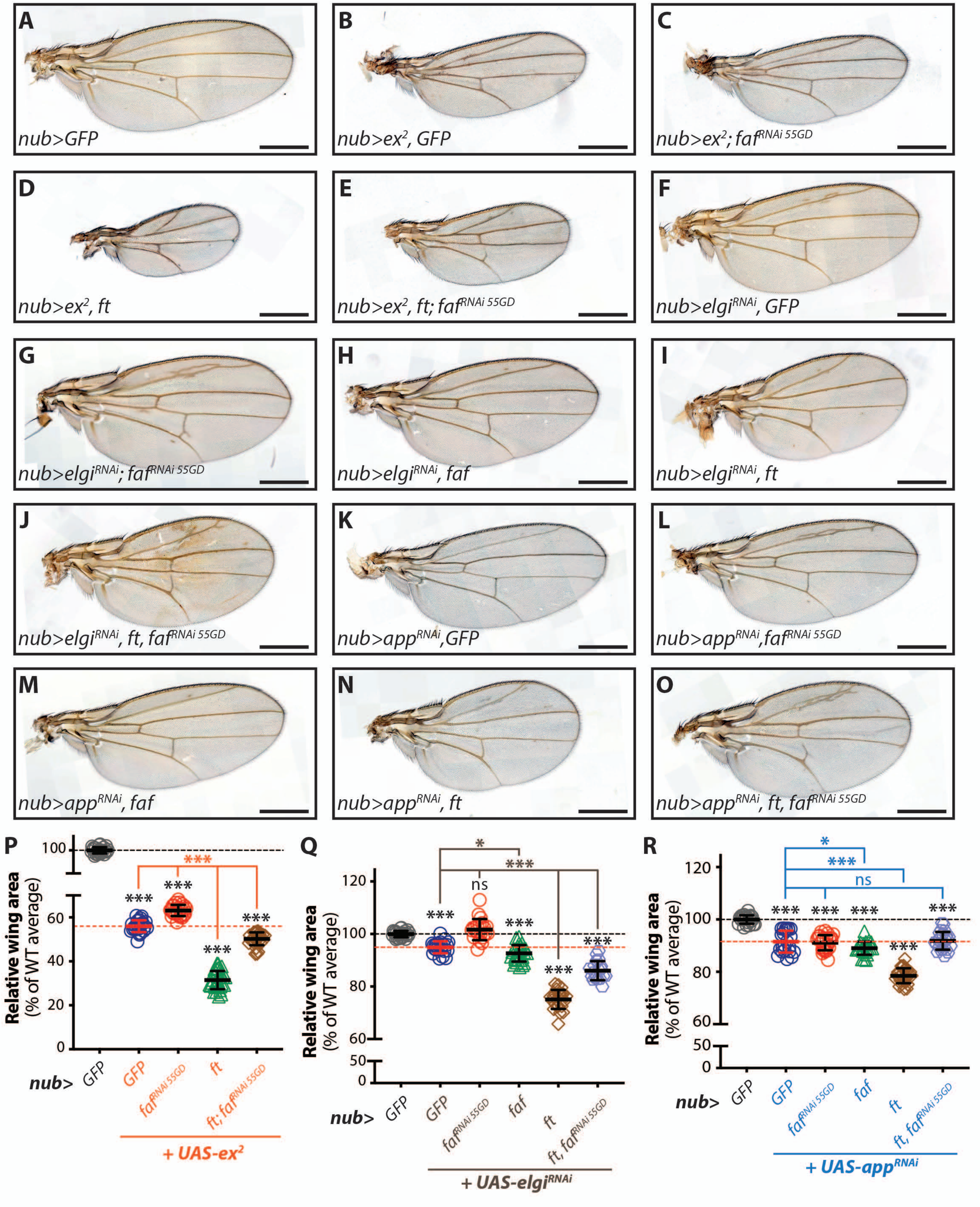


**Dawson et al. - Supplementary Figure 3**

### Supplemental Figure S4

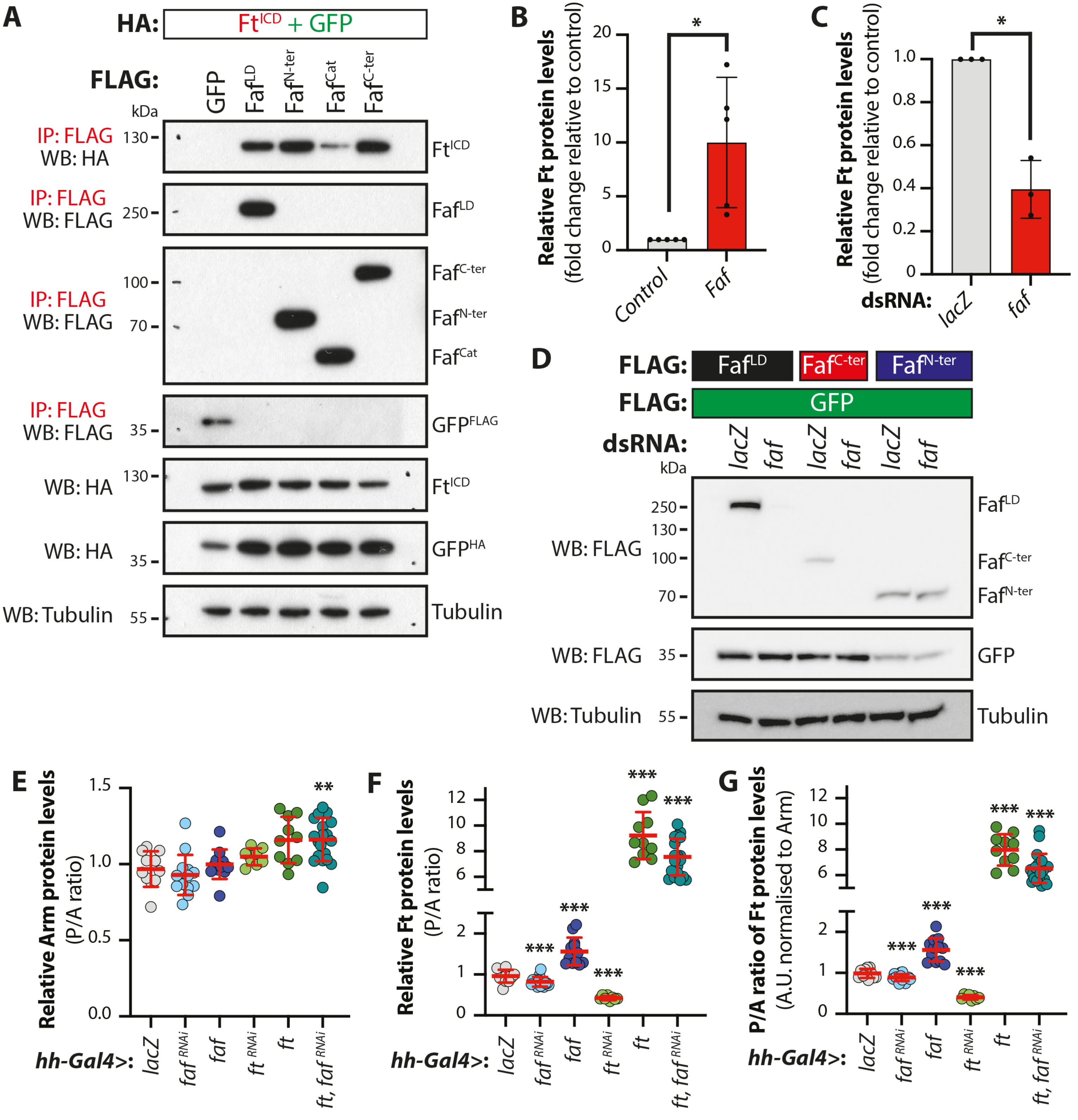


**Dawson et al. - Supplementary Figure 4**

### Supplemental Figure S5

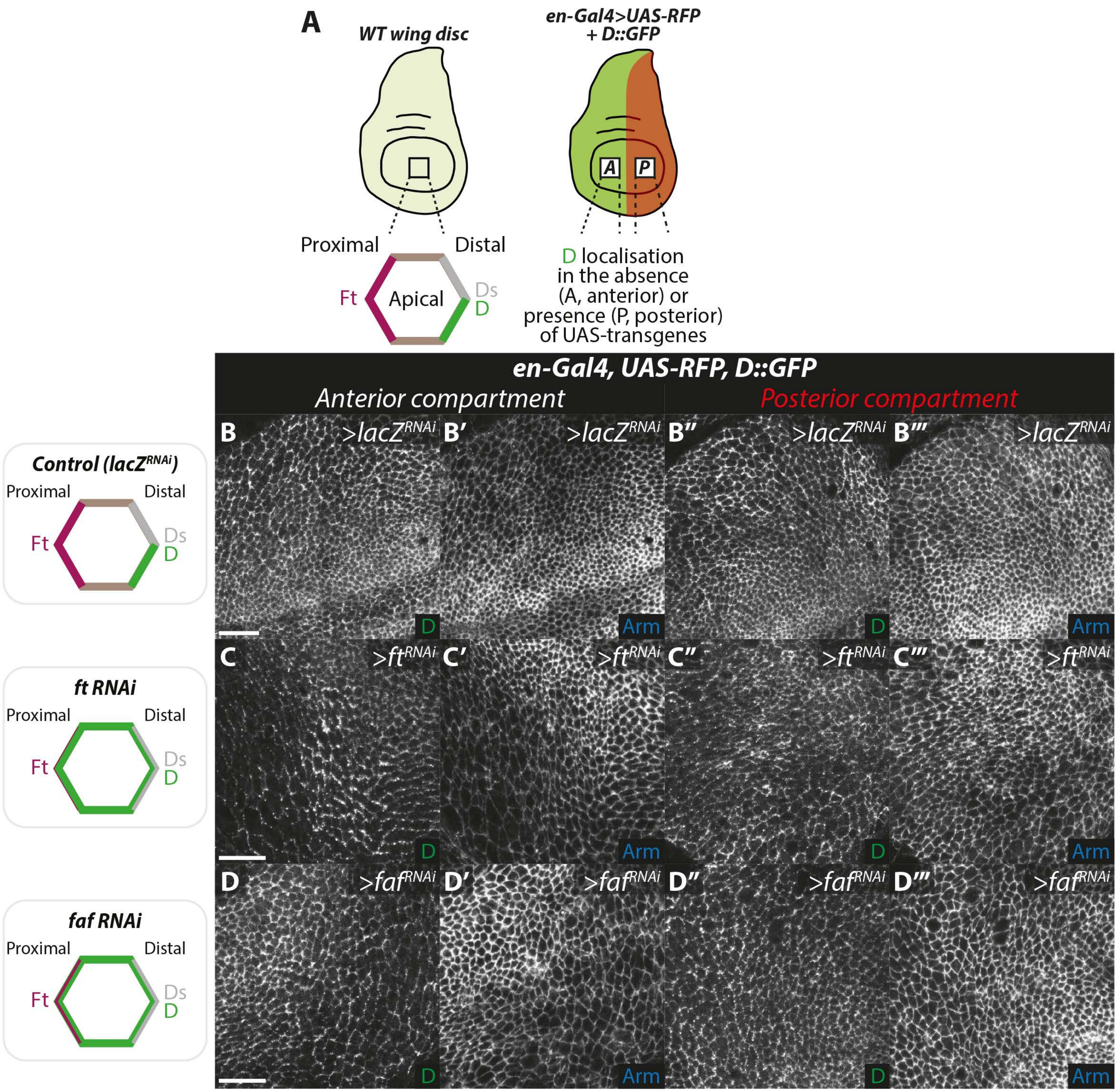


**Dawson et al. - Supplementary Figure 5**

### Supplemental Figure S6

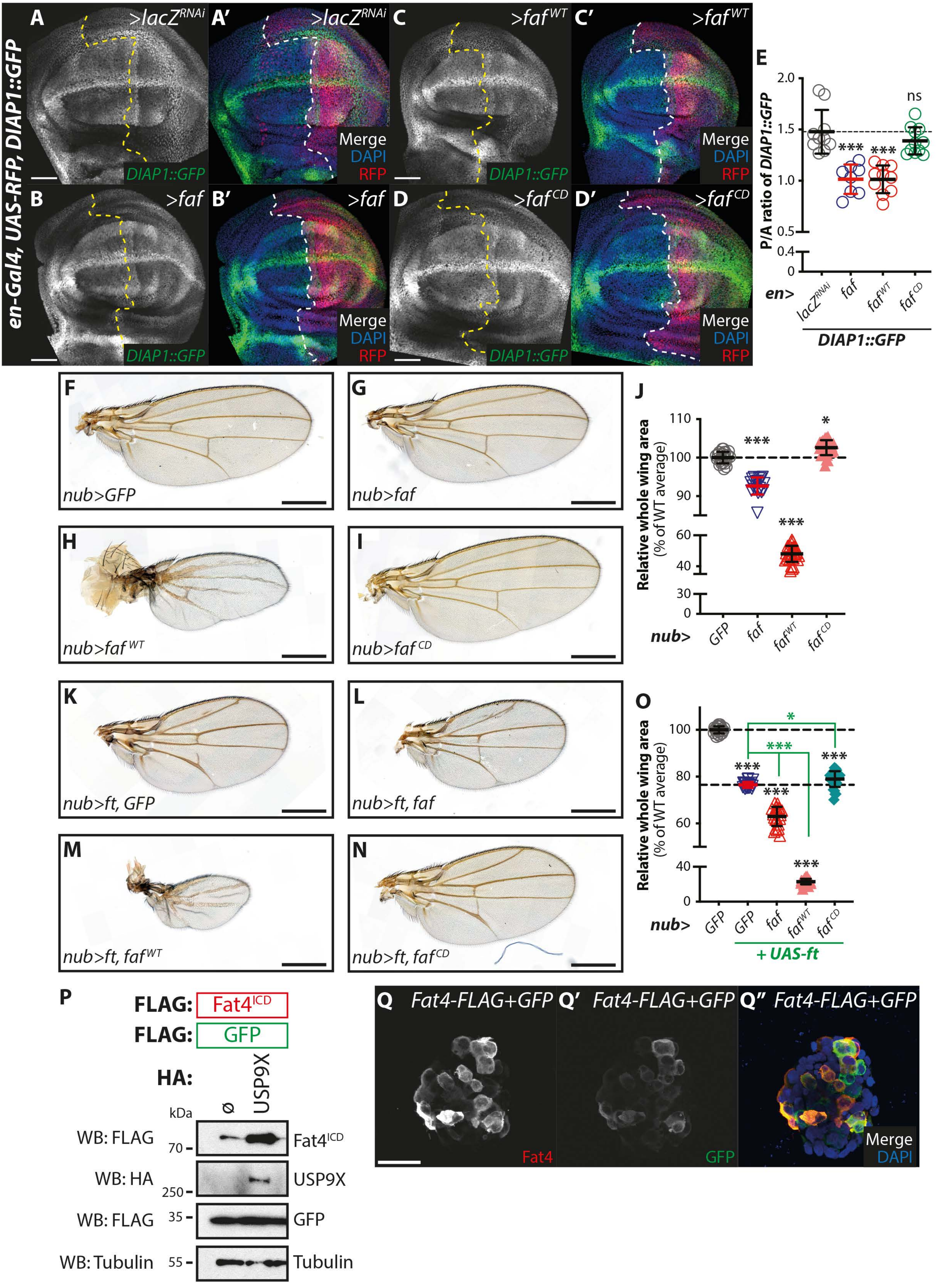


**Dawson et al. - Supplementary Figure 6**
